# An orphan *cbb*_3_-type cytochrome oxidase subunit supports *Pseudomonas aeruginosa* biofilm growth and virulence

**DOI:** 10.1101/171538

**Authors:** Jeanyoung Jo, Krista L. Cortez, William C. Cornell, Alexa Price-Whelan, Lars E.P. Dietrich

**Affiliations:** Department of Biological Sciences, Columbia University, New York, NY USA

## Abstract

Hypoxia is a common challenge faced by bacteria during associations with hosts due in part to the formation of densely packed communities (biofilms). *cbb*_3_-type cytochrome *c* oxidases, which catalyze the terminal step in respiration and have a high affinity for oxygen, have been linked to bacterial pathogenesis. The pseudomonads are unusual in that they often contain multiple full and partial (i.e., “orphan”) operons for *cbb*_3_-type oxidases and oxidase subunits. Here, we describe a unique role for the orphan catalytic subunit CcoN4 in colony biofilm development and respiration in the opportunistic pathogen *P. aeruginosa* PA14. We also show that CcoN4 contributes to the reduction of phenazines, antibiotics that support redox balancing for cells in biofilms, and to virulence in a *Caenorhabditis elegans* model of infection. These results highlight the relevance of the colony biofilm model to pathogenicity and underscore the potential of *cbb*3-type oxidases as therapeutic targets.

## INTRODUCTION

Among the oxidants available for biological reduction, molecular oxygen (O_2_) provides the highest free energy yield. Since the accumulation of O_2_ in the atmosphere between ~2.4-0.54 billion years ago (Kirschvink & Kopp 2008; Dietrich, Tice, et al. 2006), organisms that can use it for growth and survival, and tolerate its harmful byproducts, have evolved to exploit this energy and increased in complexity (Knoll & Sperling 2014; Falkowski 2006). At small scales and in crowded environments, rapid consumption of O_2_ leads to competition for this resource and has promoted diversification of bacterial and archaeal mechanisms for O_2_ reduction that has not occurred in eukaryotes (Brochier-Armanet et al. 2009). The various enzymes that allow bacteria to respire O_2_ exhibit a range of affinities and proton-pumping efficiencies and likely contribute to competitive success in hypoxic niches (Morris & Schmidt 2013). Such environments include the tissues of animal and plant hosts that are colonized by bacteria of high agricultural (Preisig et al. 1996) and clinical (Way et al. 1999; Weingarten et al. 2008) significance.

The opportunistic pathogen *Pseudomonas aeruginosa*, a colonizer of both plant and animal hosts (Rahme et al. 1995), has a branched respiratory chain with the potential to reduce O_2_ to water using at least five different terminal oxidase complexes: two quinol oxidases (*bo*_3_ and a *bd*-type cyanide insensitive oxidase) and three cytochrome *c* oxidases (*aa*_3_, *cbb*_3_-1, and *cbb*_3_-2) (**Figure 1A**). Several key publications have described *P. aeruginosa’s* complement of terminal oxidases and oxidase subunits, revealing features specific to this organism (Williams et al. 2007; Comolli & Donohue 2004; Alvarez-Ortega & Harwood 2007; Arai et al. 2014; Kawakami et al. 2010; Jo et al. 2014). *P. aeruginosa* is somewhat unusual in that it encodes two oxidases belonging to the *cbb*_3_-type family. These enzymes are notable for their relatively high catalytic activity at low O_2_ concentrations and restriction to the bacterial domain (Brochier-Armanet et al. 2009; Pitcher & Watmough 2004). (The *P. aeruginosa cbb*_3_-type oxidases are often referred to as *cbb*_3_-1 and *cbb*_3_-2; however, we will use “Cco1” and “Cco2” for these enzymes, consistent with the annotations of their encoding genes.) Most bacterial genomes that encode *cbb*_3_-type oxidases contain only one operon for such a complex, which is induced specifically under conditions of O_2_ limitation (Cosseau & Batut 2004). In *P. aeruginosa*, the *cco2* operon is induced during growth at low O_2_ concentrations, but the *cco1* operon is expressed constitutively at high levels (Comolli & Donohue 2004; Kawakami et al. 2010).

**Figure 1.**
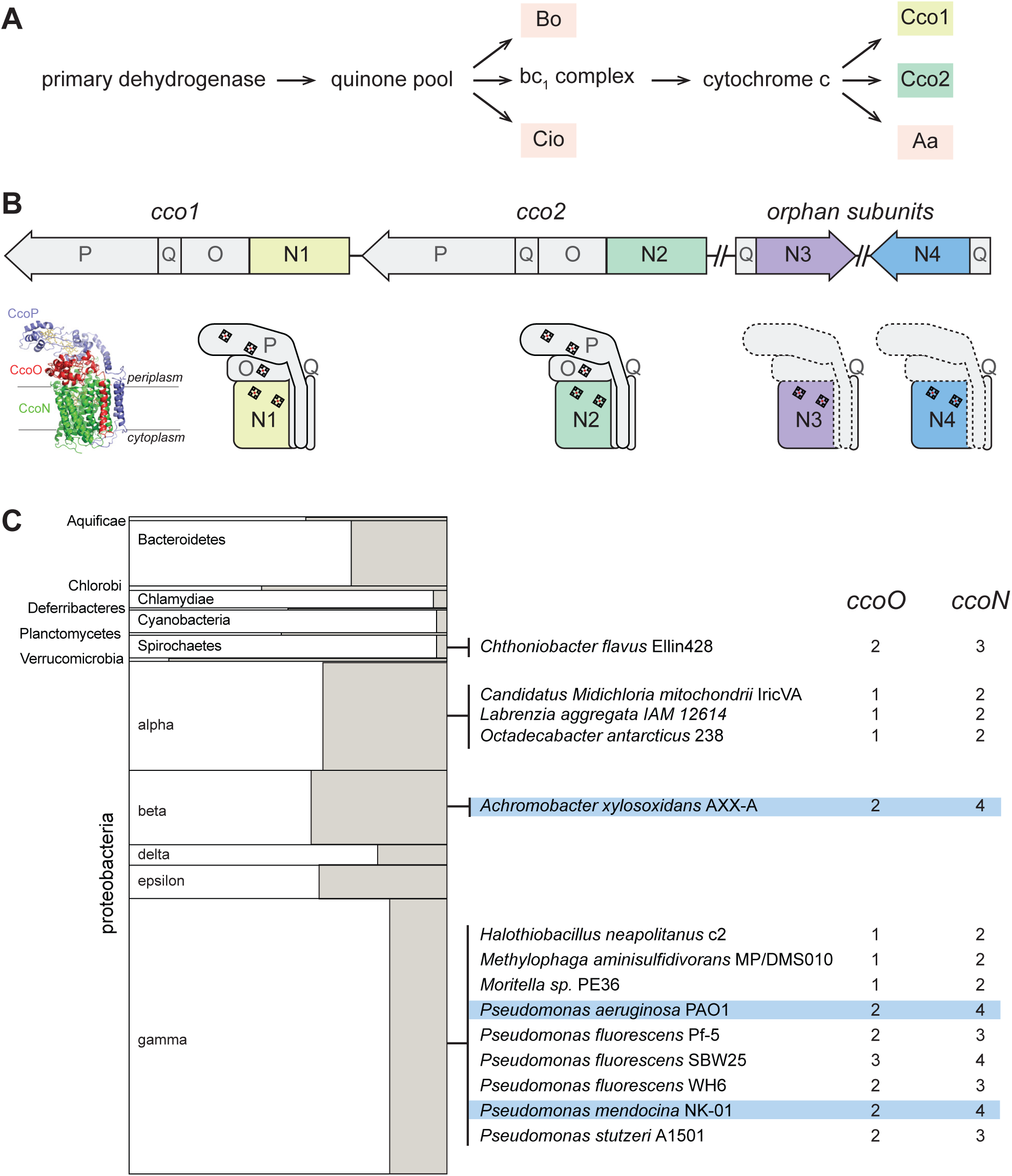
The respiratory chain and arrangement of cco genes and protein products in *P. aeruginosa*, and the phylogenetic distribution of orphan ccoN genes. (A) Branched electron transport chain in *P. aeruginosa*, containing five terminal oxidases. (B) Organization of cco genes in the *P. aeruginosa* genome. Cartoons of Cco1, Coo2 and the two orphan complexes are based on the Cco structure from *P. stutzeri* (PDB: 3mk7) (Buschmann et al. 2010). (C) Left: graphical representation of the portion of genomes in each bacterial phylum that contain *ccoO* and *N* homologs. The clades Chrysiogenetes, Gemmatimonadetes, and Zetaproteobacteria were omitted because they each contain only one species with *ccoO* and *N* homologs. The height of each rectangle indicates the total number of genomes included in the analysis. The width of each shaded rectangle represents the portion of genomes that contain *ccoN* homologs. Middle: genomes that contain more *ccoN* than *ccoO* homologs (indicating the presence of orphan *ccoN* genes) are listed. Right: numbers of *ccoO* and *ccoN* homologs in each genome. Blue highlights genomes containing more than one orphan *ccoN* homolog.

An additional quirk of the *P. aeruginosa* terminal oxidase complement lies in the presence of genes for “orphan” *cbb*_3_-type subunits at chromosomal locations distinct from the *cco1* and *cco2* operons. While the *cco1* and *cco2* operons, which are chromosomally adjacent, each contain four genes encoding a functional Cco complex (consisting of subunits N, O, P, and Q), the two additional partial operons *ccoN3Q3* and *ccoN4Q4* each contain homologs coding for only the Q and catalytic N subunits (**Figure 1B**). Expression of the *ccoN3Q3* operon is induced under anaerobic denitrification conditions (Alvarez-Ortega & Harwood 2007), and by nitrite exposure during growth under 2% O_2_ (Hirai et al. 2016). During aerobic growth in liquid cultures, *ccoN4Q4* is induced by cyanide, which is produced in stationary phase (Hirai et al. 2016). However, additional expression studies indicate that *ccoN4Q4* transcription is influenced by redox conditions, as this operon is induced by O_2_ limitation (Alvarez-Ortega & Harwood 2007) and slightly downregulated in response to pyocyanin, a redox-active antibiotic produced by *P. aeruginosa* (Dietrich, Price-Whelan, et al. 2006).

In a recent study, Hirai et al. characterized the biochemical properties and physiological roles of *P. aeruginosa *cbb*_3_* isoforms containing combinations of canonical and orphan subunits (Hirai et al. 2016). In a strain lacking all of the aerobic terminal oxidases, expression of any isoform conferred the ability to grow using O_2_, confirming that isoforms containing the orphan N subunits are functional. Furthermore, the authors found that the products of *ccoN3Q3* and *ccoN4Q4* contributed resistance to nitrite and cyanide, respectively, during growth in liquid cultures under low-O_2_ conditions. While these results provide insight into contributions of the *cbb*_3_ heterocomplexes to growth in liquid cultures, potential roles for N3- and N4-containing isoforms in biofilm growth and pathogenicity have yet to be explored.

The biofilm lifestyle—in which cells grow in a dense community encased in a self-produced matrix—has been linked to the establishment and persistence of infections in diverse systems (Edwards & Kjellerup 2012; Rybtke et al. 2015). Biofilm development promotes the formation of O_2_ gradients such that cells at a distance from the biofilm surface are subjected to hypoxic or anoxic conditions (Werner et al. 2004). We have shown that O_2_ limitation for cells in biofilms leads to an imbalance in the intracellular redox state. This can be relieved by a change in community morphology, which increases the surface area-to-volume ratio of the biofilm and therefore access to O_2_ for resident cells (Kempes et al. 2014). For *P. aeruginosa* cells in biofilms, the intracellular accumulation of reducing power can also be prevented by production and reduction of endogenous antibiotics called phenazines, which mediate extracellular electron transfer to oxidants available at a distance (Dietrich et al. 2013). We have found that biofilm-specific phenazine production contributes to pathogenicity in a murine model of acute pulmonary infection (Recinos et al. 2012), further underscoring the importance of phenazine-mediated redox balancing for *P. aeruginosa* cells in communities.

Because of the formation of an O_2_ gradient inherent to the biofilm lifestyle, we hypothesized that the differential regulation of the *P. aeruginosa cco* operons affects their contributions to metabolic electron flow in biofilm subzones. We evaluated the roles of various *cbb*_3_-type oxidase isoforms in multicellular behavior and virulence. Our results indicate that isoforms containing the orphan subunit CcoN4 can support survival in biofilms via O_2_ and phenazine reduction and contribute to *P. aeruginosa* pathogenicity in a *Caenorhabditis elegans* “slow killing” model of infection.

## RESULTS

### A small minority of bacterial genomes encode cbb_3_-type oxidase subunits in partial (“orphan”) operons

Biochemical, genetic, and genomic analyses suggest that the CcoN and CcoO subunits, typically encoded by an operon, form the minimal functional unit of *cbb*_3_-type oxidases (Ducluzeau et al. 2008; de Gier et al. 1996; Zufferey et al. 1996). CcoN is the membrane-integrated catalytic subunit and contains two *b*-type hemes and a copper ion. CcoO is membrane-anchored and contains one *c*-type heme. Additional redox subunits and/or subunits implicated in complex assembly, such as CcoQ and CcoP, can be encoded by adjacent genes (**Figure 1B**). *ccoNO*-containing clusters are widely distributed across phyla of the bacterial domain (Ducluzeau et al. 2008). We used the EggNOG database, which contains representative genomes for more than 3000 bacterial species (Huerta-Cepas et al. 2016) to obtain an overview of the presence and frequency of *cco* genes. Out of 3318 queried bacterial genomes we found 467 with full *cco* operons (encoding potentially functional *cbb*_3_-type oxidases with O and N subunits). Among these, 78 contain more than one full operon. We also used EggNOG to look for orphan *ccoN* genes by examining the relative numbers of *ccoO* and *ccoN* homologs in individual genomes. We found 14 genomes, among which *Pseudomonas* species are overrepresented, that contain orphan *ccoN* genes (**Figure 1C**), and our analysis yielded 3 species that contain more than one orphan *ccoN* gene: *Pseudomonas mendocina*, *Pseudomonas aeruginosa*, and *Achromobacter xylosoxidans*. *P. mendocina* is a soil bacterium and occasional nosocomial pathogen that is closely related to *P. aeruginosa*, based on 16S rRNA gene sequence comparison (Anzai et al. 2000). *A. xylosoxidans*, in contrast, is a member of a different proteobacterial class but nevertheless is often mistaken for *P. aeruginosa* (Saiman et al. 2001). Like *P. aeruginosa*, it is an opportunistic pathogen that can cause pulmonary infections in immunocompromised individuals and patients with cystic fibrosis (De Baets et al. 2007; Firmida et al. 2016). Hirai et al. previously reported a ClustalW-based analysis of CcoN homologs specifically from pseudomonads, which indicated the presence of orphan genes in additional species not represented in the EggNOG database. These include *P. denitrificans*, which contains two orphan genes (Hirai et al. 2016).

### CcoN4-containing isoforms function specifically in biofilms to support community morphogenesis and respiration

During growth in a biofilm, subpopulations of cells are subjected to regimes of electron donor and O_2_ availability that may create unique metabolic demands and require modulation of the respiratory chain for survival (Alvarez-Ortega & Harwood 2007; Borriello et al. 2004; Werner et al. 2004). We therefore investigated the contributions of individual *cco* genes and gene clusters to *P. aeruginosa* PA14 biofilm development using a colony morphology assay, which has demonstrated sensitivity to electron acceptor availability and utilization (Dietrich et al. 2013). Because the Cco1 and Cco2 complexes are the most important cytochrome oxidases for growth of *P. aeruginosa* in fully aerated and O_2_-limited liquid cultures (Alvarez-Ortega & Harwood 2007; Arai et al. 2014), we predicted that mutations disabling the functions of Cco1 and Cco2 would affect colony growth. Indeed, a mutant lacking both the *cco1* and *cco2* operons (“Δ*cco1cco2*”) produced thin biofilms with a smaller diameter than the wild type. After five days of development, this mutant displayed a dramatic phenotype consisting of a tall central ring feature surrounded by short ridges that emanate radially (**Figure 2A**; **Figure 2— figure supplement 1**). Δ*cco1cco2* colonies were also darker in color, indicating increased uptake of the dye Congo red, which binds to the extracellular matrix produced by biofilms (Friedman & Kolter 2004). Surprisingly, a strain specifically lacking the catalytic subunits of Cco1 and Cco2 (“Δ*N1*Δ*N2*”), while showing a growth defect similar to that of Δ*cco1cco2* when grown in liquid culture (**Figure 2C**), showed biofilm development that was similar to that of the wild type (**Figure 2A**; **Figure 2‐‐figure supplement 1**). As it is known that CcoN3 and CcoN4 can form functional complexes with subunits of the Cco1 and Cco2 oxidases in *P. aeruginosa* PAO1 (Hirai et al. 2016), this led us to hypothesize that Cco isoforms containing the orphan subunits CcoN3 and/or CcoN4 could substitute for Cco1 and Cco2 in the biofilm context. Deleting *ccoN3* (“Δ*N3*” or “Δ*N1*Δ*N2*Δ*N3*”) did not have an observable effect on biofilm development when mutants were compared to respective parent strains (**Figure 2— figure supplement 1**). However, the phenotype of a “Δ*N1*Δ*N2*Δ*N4*” mutant was consistent with our model, as it mimicked that of the Δ*cco1cco2* mutant in both liquid-culture and biofilm growth (**Figures 2A and 2C; Figure 2— figure supplement 1**). Furthermore, we found that a mutant lacking only *ccoN4* (“Δ*N4*”) displayed an altered phenotype in that it began to form wrinkle structures earlier than the wild type (**Figure 2— figure supplement 1**), which developed into a disordered region of wrinkles inside a central ring, surrounded by long, radially emanating ridges (**Figure 2A**). Reintroduction of the *ccoN4* gene into either of these strains restored the phenotypes of the respective parent strains (**Figure 2— figure supplement 1**). The developmental pattern of the Δ*N4* colony is reminiscent of those displayed by mutants defective in phenazine production and sensing (**Figure 2— figure supplement 1**) (Dietrich et al. 2008; Dietrich et al. 2013; Sakhtah et al. 2016; Okegbe et al. 2017). Although Δ*N4* itself showed a unique phenotype in the colony morphology assay, its growth in shaken liquid cultures was indistinguishable from that of the wild type (**Figure 2C**). These results suggest that CcoN4-containing Cco isoform(s) play physiological roles that are specific to the growth conditions encountered in biofilms.

**Figure 2.**
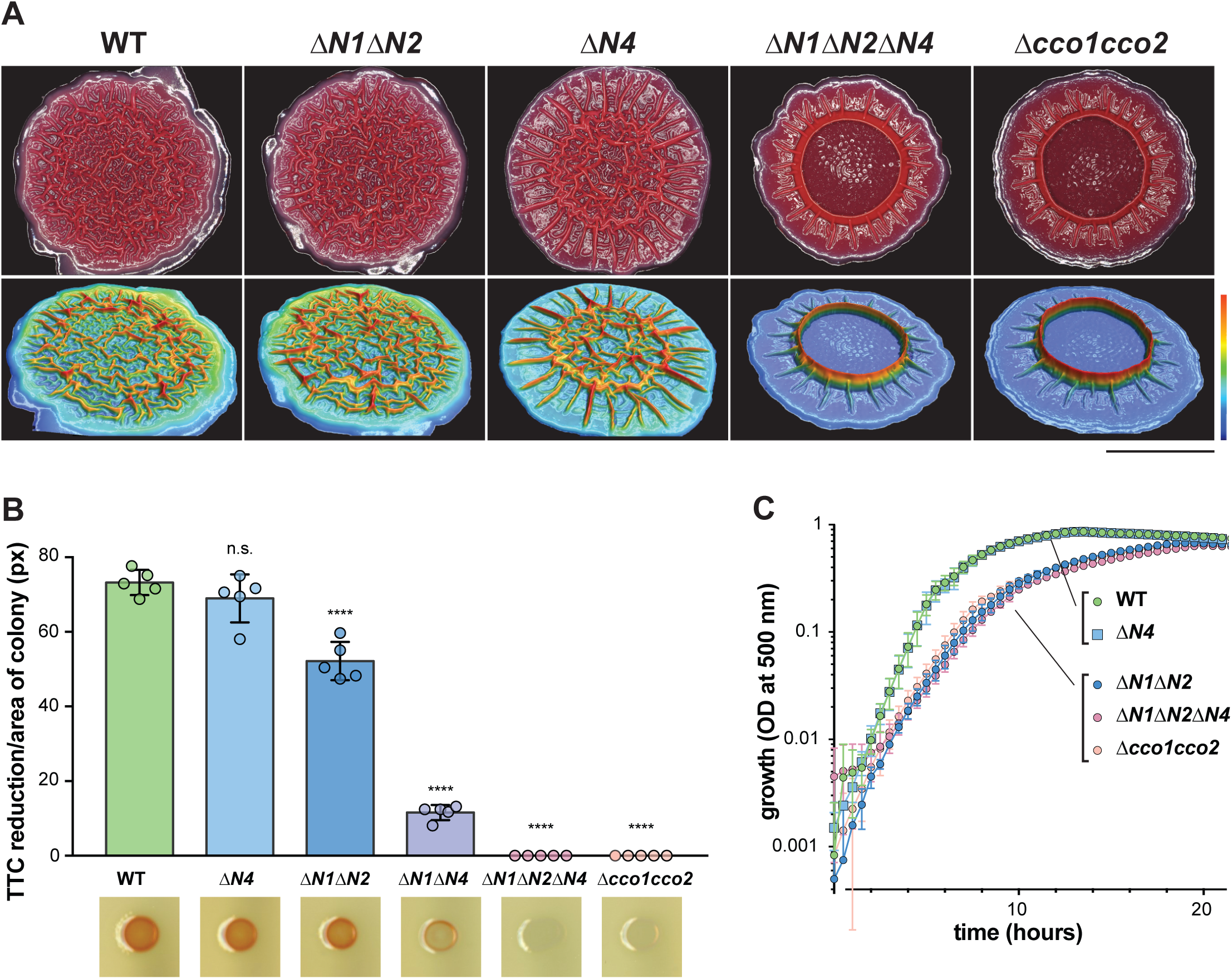
CcoN4-containing heterocomplexes make biofilm-specific contributions to morphogenesis and respiration. (A) Top: Five-day-old colony biofilms of PA14 wild type and *cco* mutant strains. Biofilm morphologies are representative of more than ten biological replicates. Images were generated using a Keyence digital microscope. Scale bar is 1 cm. Bottom: 3D surface images of the biofilms shown in the top panel. Images were generated using a Keyence wide-area 3D measurement system. Height scale bar: bottom (blue) to top (red) is 0 - 0.7 mm for WT, Δ*N1*Δ*N2*, and Δ*N4*; 0 - 1.5 mm for Δ*N1*Δ*N2*Δ*N4* and Δ*cco1cco2.* (B) TTC reduction by *cco* mutant colonies after one day of growth. Upon reduction, TTC undergoes an irreversible color change from colorless to red. Bars represent the average, and error bars represent the standard deviation, of individually plotted biological replicates (n = 5). P-values were calculated using unpaired, two-tailed t tests comparing each mutant to wild type (^∗∗∗^, P ≤ 0.001; ^∗∗∗∗^, P ≤ 0.0001). For full statistical reporting, refer to Table 4. (C) Mean growth of PA14 wild type and *cco* mutant strains in MOPS defined medium with 20 mM succinate. Error bars represent the standard deviation of biological triplicates.

Next, we asked whether CcoN4 contributes to respiration in biofilms. We tested a suite of *cco* mutants for reduction of triphenyl tetrazolium chloride (TTC), an activity that is often associated with cytochrome *c* oxidase-dependent respiration (Rich et al. 2001). The Δ*cco1cco2* mutant showed a severe defect in TTC reduction, which was recapitulated by the Δ*N1*Δ*N2*Δ*N4* mutant. As in the colony morphology assay, this extreme phenotype was not recapitulated in a mutant lacking only CcoN1 and CcoN2, indicating that CcoN4 contributes to respiratory activity in PA14 biofilms. Although we did not detect a defect in TTC reduction for the Δ*N4* mutant, we saw an intermediate level of TTC reduction for a “Δ*N1*Δ*N4*” mutant compared to Δ*N1*Δ*N2* and Δ*N1*Δ*N2*Δ*N4*, further implicating the CcoN4 subunit in this activity (**Figure 2B**).

A recent study implicated CcoN4 in resistance to cyanide, a respiratory toxin that is produced by *P. aeruginosa* (Hirai et al. 2016). The altered biofilm phenotypes of Δ*N4* mutants could therefore be attributed to an increased sensitivity to cyanide produced during biofilm growth. We deleted the *hcn* operon, coding for cyanide biosynthetic enzymes, in the wild-type, phenazine-null, and various *cco* mutant backgrounds. The biofilm morphologies and liquid-culture growth of these strains were unaffected by the Δ*hcnABC* mutation, indicating that the biofilm-specific role of CcoN4 explored in this work is independent of its role in mediating cyanide resistance (**Figure 2— figure supplement 2**). Additionally, we examined genomes available in the Pseudomonas Genome Database for the presence of homologs encoding CcoN subunits (*ccoN* genes) and enzymes for cyanide synthesis (*hcnABC*) (Winsor et al. 2016) and did not find a clear correlation between the presence of *hcnABC* and *ccoN4* homologs (**Figure 2— figure supplement 3**).

Together, the effects of *cco* gene mutations that we observed in assays for colony morphogenesis and TTC reduction suggest that one or more CcoN4-containing Cco isoform(s) support respiration and redox balancing, and is/are utilized preferentially in comparison to CcoN1- and CcoN2-containing Cco complexes, in biofilms.

### Different CcoN subunits are required for competitive fitness in early or late colony development

To further test CcoN4’s contribution to growth in biofilms, we performed competition assays in which Δ*N4* and other mutants were grown as mixed-strain biofilms with the wild type. In each of these assays, one strain was labeled with constitutively-expressed YFP so that the strains could be distinguished during enumeration of colony forming units (CFUs). Experiments were performed with the label on each strain to confirm that YFP expression did not affect fitness (**Figure 3— figure supplement 1A, B**). When competitive fitness was assessed after three days of colony growth (**Figure 3A**), Δ*N4* cells showed a disadvantage, with the wild type outcompeting Δ*N4* by a factor of two. This was similar to the disadvantage observed for the Δ*N1*Δ*N2* mutant, further suggesting that the orphan subunit CcoN4 plays a significant role in biofilm metabolism. Remarkably, deletion of *ccoN4* in mutants already lacking *ccoN1* and *ccoN2* led to a drastic decrease in fitness, with the wild type outcompeting Δ*N1*Δ*N2*Δ*N4* by a factor of 16. This disadvantage was comparable to that observed for the mutant lacking the full *cco* operons (Δ*cco1cco2*), underscoring the importance of CcoN4-containing isoforms during biofilm growth.

**Figure 3.**
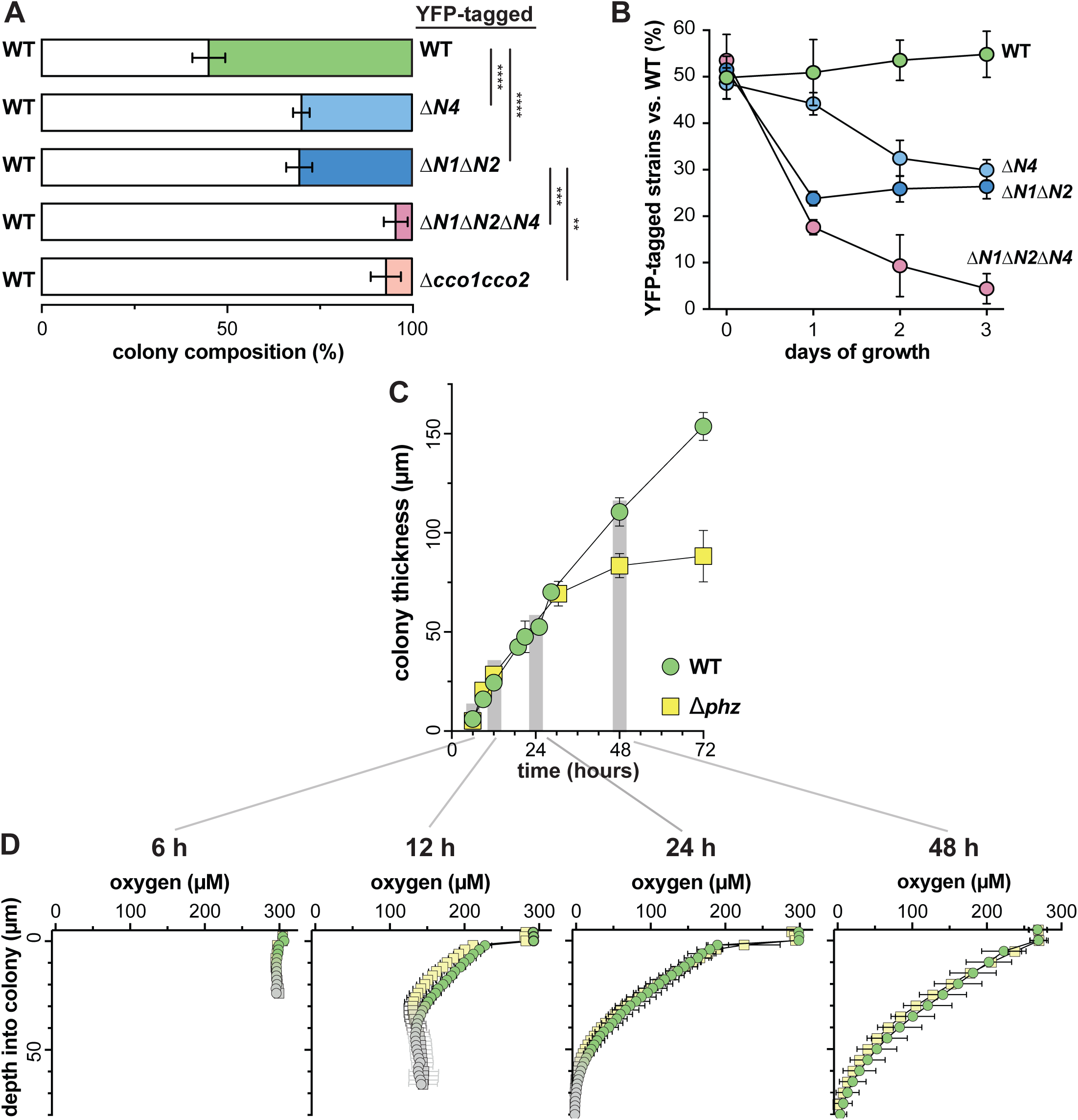
CcoN4 is necessary for optimal fitness in biofilms, particularly when oxygen becomes limiting. (A) Relative fitness of various YFP-labeled *cco* mutants when co-cultured with WT in mixed-strain biofilms for three days. Error bars represent the standard deviation of biological triplicates. P-values were calculated using unpaired, two-tailed t tests (^∗∗∗^, P ≤ 0.001; ^∗∗∗∗^, P ≤ 0.0001). For full statistical reporting, refer to Table 4. (B) Time course showing relative fitness, over a period of three days, of various *cco* mutants when co-cultured with WT in mixed-strain biofilms. Results are shown for experiments in which the WT was co-cultured with various “labeled” strains, i.e. those that were engineered to constitutively express YFP (See Figure 3— figure supplement 1 for results from experiments in which the labeled WT was co-cultured with unlabeled mutants.) Error bars represent the standard deviation of biological triplicates. (C) Change in thickness over three days of development for colony biofilms of WT and Δ*phz* as assessed by thin sectioning and DIC microscopy. After the onset of wrinkling, thickness was determined for the base (i.e., the “valley” between wrinkles). Error bars represent the standard deviation of biological triplicates. (D) O_2_ profiles of colonies at selected timepoints within the first three days of biofilm development. Green, WT; yellow, *Δ*phz*;* gray, outside the colony (measurements made in the agar directly below the colony). Error bars denote standard deviation of biological triplicates.

To further explore the temporal dynamics of N subunit utilization, we repeated the competition assay, but sampled each day over the course of three days (**Figure 3B**). The fitness disadvantage that we had found for strains lacking CcoN1 and CcoN2 was evident after only one day of growth and did not significantly change after that. In contrast, the Δ*N4*-specific decline in fitness did not occur before the second day. These data suggest that the contributions of the various N subunits to biofilm metabolism differ depending on developmental stage.

DIC imaging of thin sections from wild-type colonies reveals morphological variation over depth that may result from decreasing O_2_ availability (**Figure 3— figure supplement 1C**). We have previously reported that three-day-old PA14 colony biofilms are hypoxic at depth (Dietrich et al. 2013) and that O_2_ availability is generally higher in thinner biofilms, such as those formed by a phenazine-null mutant (Δ*phz*). We have proposed that the utilization of phenazines as electron acceptors in wild-type biofilms enables cellular survival in the hypoxic zone and promotes colony growth (Okegbe et al. 2014). The relatively late-onset phenotype of the Δ*N4* mutant in the competition assay suggested to us that CcoN4 may play a role in survival during formation of the hypoxic colony subzone and that this zone could arise at a point between one and two days of colony growth. We measured O_2_ concentrations in wild-type and Δ*phz* biofilms at specific time points over development, and found that O_2_ declined similarly with depth in both strains (**Figure 3D**). The rate of increase in height of Δ*phz* tapered off when a hypoxic zone began to form, consistent with our model that the base does not increase in thickness when electron acceptors (O_2_ or phenazines) are not available. Although we cannot pinpoint the exact depth at which the O_2_ microsensor leaves the colony base and enters the underlying agar, we can estimate these values based on colony thickness measurements (**Figure 3C**). When we measured the thickness of wild-type and Δ*phz* biofilms over three days of incubation, we found that the values began to diverge between 30 and 48 hours of growth, after the colonies reached ~70 μm in height, which coincides with the depth at which O_2_ becomes undetectable. Δ*phz* colonies reached a maximum thickness of ~80 μm, while wild-type colonies continued to grow to ~150 μm (**Figure 3C**). In this context, it is interesting to note that the point of divergence for the increase in wild-type and Δ*phz* colony thickness corresponds to the point at which CcoN4 becomes important for cell viability in our mixed-strain colony growth experiments (**Figure 3B**). We hypothesize that this threshold thickness leads to a level of O_2_ limitation that is physiologically relevant for the roles of phenazines and CcoN4 in biofilm metabolism.

### cco genes show differential expression across biofilm subzones

*P. aeruginosa’s* five canonical terminal oxidases are optimized to function under and in response to distinct environmental conditions, including various levels of O_2_ availability (Arai et al. 2014; Kawakami et al. 2010; Alvarez-Ortega & Harwood 2007; Comolli & Donohue 2004). Furthermore, recent studies, along with our results, suggest that even within the Cco terminal oxidase complexes, the various N subunits may perform different functions (Hirai et al. 2016). We sought to determine whether differential regulation of *cco* genes could lead to uneven expression across biofilm subzones. To test this, we engineered reporter strains in which GFP expression is regulated by the *cco1*, *cco2*, or *ccoN4Q4* promoters. Biofilms of these strains were grown for three days, thin-sectioned, and imaged by fluorescence microscopy. Representative results are shown in the left panel of **Figure 4**. The right panel of **Figure 4** contains plotted GFP signal intensity and O_2_ concentration measurements over depth for PA14 wild-type colonies. *cco1* and *ccoN4* expression patterns indicate that the Cco1 oxidase and the CcoN4 subunit are produced throughout the biofilm (**Figure 4**). *cco2* expression, on the other hand, is relatively low in the top portion of the biofilm and shows a sharp induction starting at a depth of ~45 μm. This observation is consistent with previous studies showing that *cco2* expression is regulated by Anr, a global transcription factor that controls gene expression in response to a shift from oxic to anoxic conditions (Comolli & Donohue 2004; Kawakami et al. 2010; Ray & Williams 1997).

**Figure 4.**
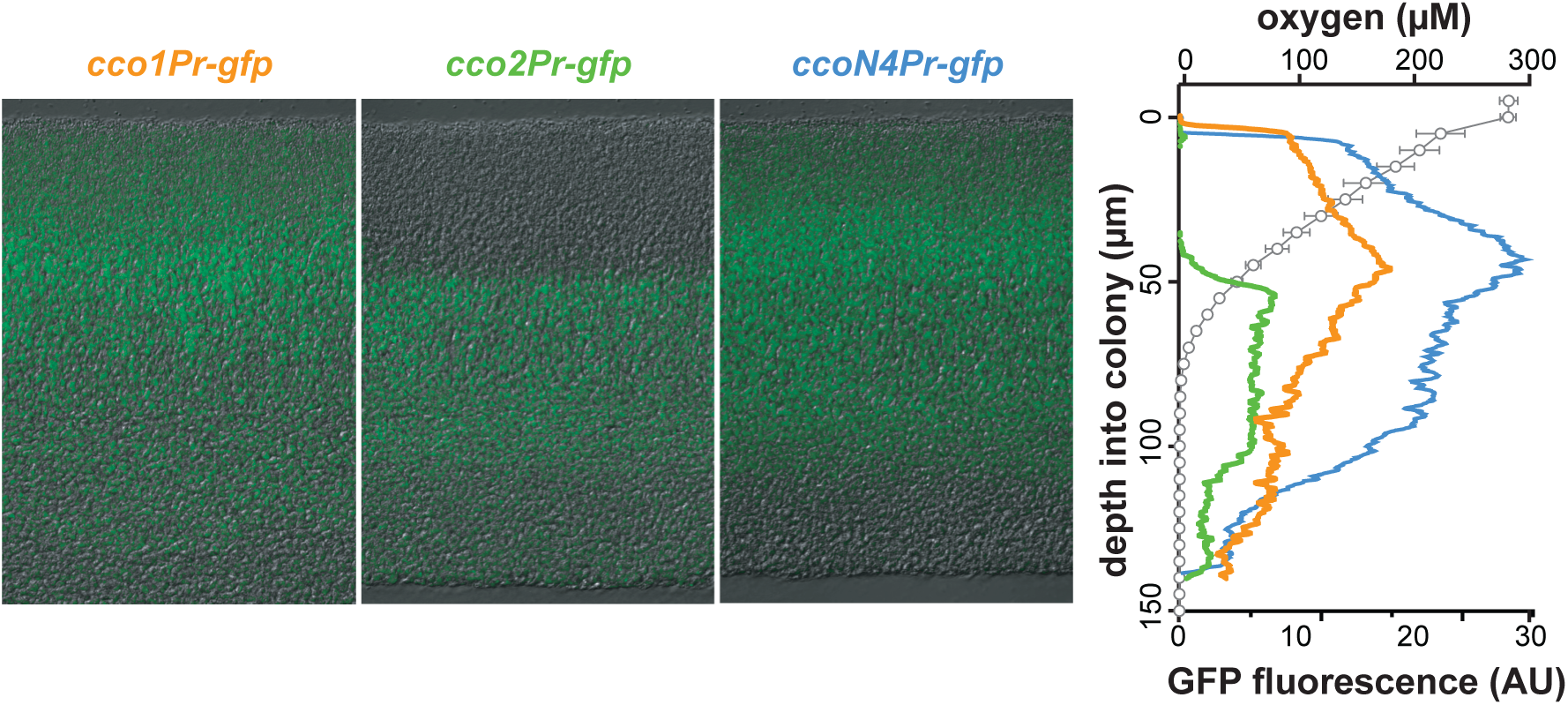
*cco* genes are differentially expressed over biofilm depth. Left: Representative images of thin sections prepared from WT biofilms grown for three days. Each biofilm is expressing a translational GFP reporter under the control of the *cco1*, *cco2*, or *ccoQ4N4* promoter. Reporter fluorescence is shown in green and overlain on respective DIC images. Right: Fluorescence values corresponding to images on the left. Fluorescence values for the empty MCS control have been subtracted from each respective plot. O_2_ concentration (open circles) over depth from three-day-old WT biofilms is also shown. Error bars represent the standard deviation of biological triplicates and are not shown in cases where they would be obscured by the point markers. y-axis in the right panel provides a scale bar for the left panel. Reporter fluorescence images and values are representative of four biological replicates.

Though previous studies have evaluated expression as a function of growth phase in shaken liquid cultures for *cco1* and *cco2*, this property has not been examined for *ccoN4Q4.* We monitored the fluorescence of our engineered *cco* gene reporter strains during growth under this condition in a nutrient-rich medium. As expected based on the known constitutive expression of *cco1* and Anr-dependence of *cco2* induction, we saw *cco1*-associated fluorescence increase before that associated with *cco2.* Induction of *ccoN4Q4* occurred after that of *cco1* and *cco2* (**Figure 4— figure supplement 1**), consistent with microarray data showing that this locus is strongly induced by O_2_ limitation (Alvarez-Ortega & Harwood 2007). However, our observation that *ccoN4Q4* is expressed in the aerobic zone, where *cco2* is not expressed, in biofilms (**Figure 4**) suggests that an Anr-independent mechanism functions to induce this operon during multicellular growth.

Our results indicate that different Cco isoforms may function in specific biofilm subzones, but that CcoN4-containing isoforms could potentially form throughout the biofilm. These data, together with our observation that Δ*N4* biofilms exhibit a fitness disadvantage from day two (**Figure 3B**), led us to more closely examine the development and chemical characteristics of the biofilm over depth.

### Microelectrode-based redox profiling reveals differential phenazine reduction activity in wild-type and cco mutant biofilms

The results shown in **Figure 2B** implicate CcoN4-containing isoforms in the reduction of TTC, a small molecule that interacts with the respiratory chain (Rich et al. 2001). Similar activities have been demonstrated for phenazines, including the synthetic compound phenazine methosulfate (Nachlas et al. 1960) and those produced naturally by *P. aeruginosa* (Armstrong & Stewart-Tull 1971). Given that CcoN4 and phenazines function to influence morphogenesis at similar stages of biofilm growth (**Figures 2A, 3, Figure 2— figure supplement 1, Figure 3— figure supplement 1A, B**), we wondered whether the role of CcoN4 in biofilm development was linked to phenazine metabolism. We used a Unisense platinum microelectrode with a 20-30 μm tip to measure the extracellular redox potential in biofilms as a function of depth. This electrode measures the inclination of the sample to donate or accept electrons relative to a Ag/AgCl reference electrode. We found that wild-type colonies showed a decrease in redox potential over depth, indicating an increased ratio of reduced to oxidized phenazines, while the redox potential of Δ*phz* colonies remained unchanged (**Figure 5A**). To confirm that phenazines are the primary determinant of the measured redox potential in the wild type, we grew Δ*phz* colonies on medium containing phenazine methosulfate (a synthetic compound that resembles the natural phenazines that regulate *P. aeruginosa* colony morphogenesis (Sakhtah et al. 2016)), and found that these colonies yielded redox profiles similar to those of the wild type (**Figure 5— figure supplement 1A**). Therefore, though the microelectrode we employed is capable of interacting with many redox-active substrates, we found that its signal was primarily determined by phenazines in our system. In addition, while wild-type colonies showed rapid decreases in O_2_ availability starting at the surface, the strongest decrease in redox potential was detected after ~50 μm (**Figure 5A**). These results suggest that the bacteria residing in the biofilm differentially utilize O_2_ and phenazines depending on their position and that O_2_ is the preferred electron acceptor.

**Figure 5.**
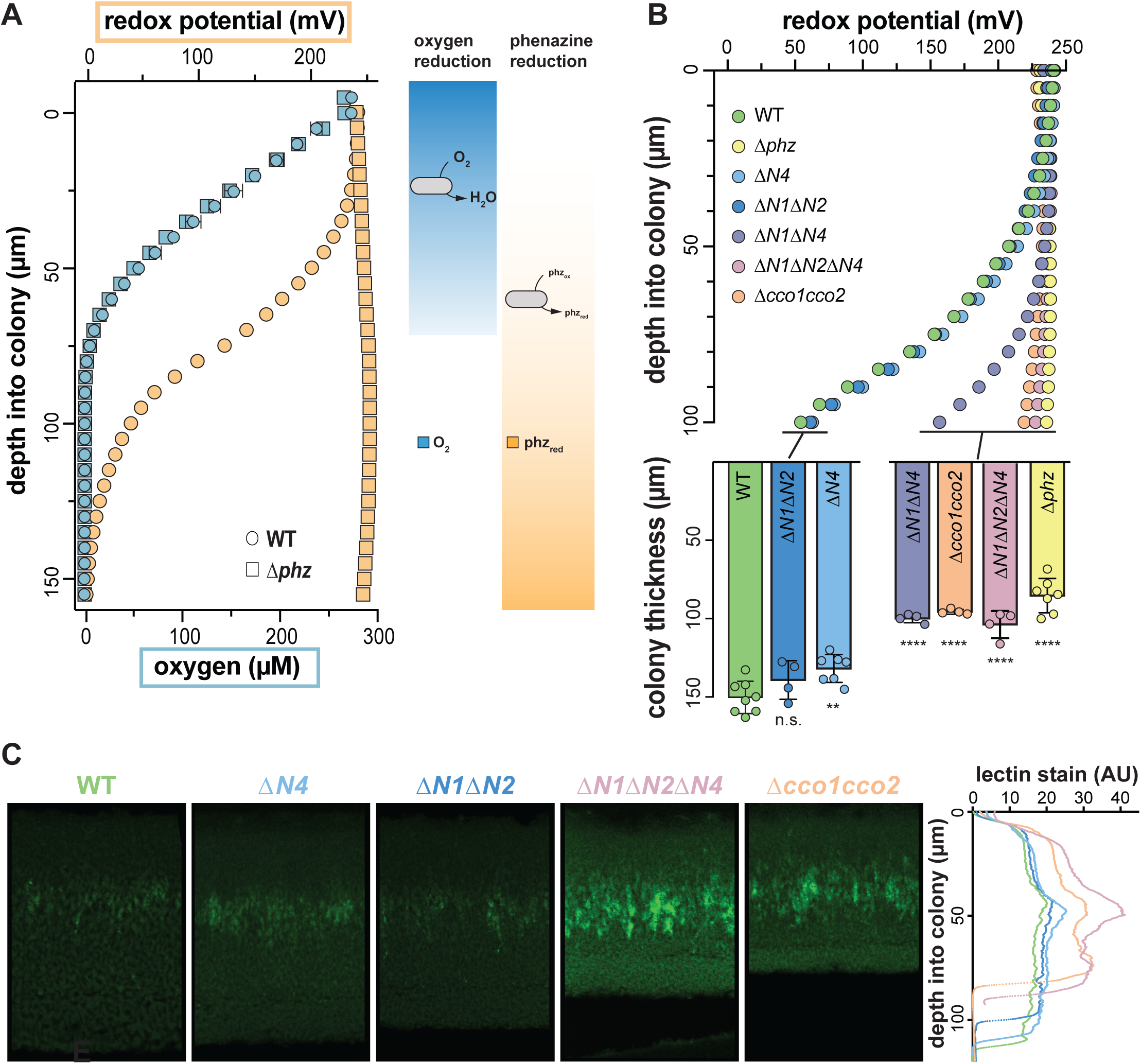
Characterization of chemical gradients and matrix distribution in PA14 WT and mutant colony biofilms. (A) Left: Change in O_2_ concentration (blue) and redox potential (orange) with depth for WT and Δ*phz* biofilms grown for two days. For O_2_ profiles, error bars represent the standard deviation of biological triplicates. For redox profiles, data are representative of at least five biological replicates. Right: model depicting the distribution of O_2_ and reduced vs. oxidized phenazines in biofilms. (B) Top: Change in redox potential with depth for WT and various mutant biofilms grown for two days. Data are representative of at least five biological replicates. Bottom: Thickness of three-day-old colony biofilms of the indicated strains. Bars represent the average of the plotted data points (each point representing one biological replicate, n ≥ 4), and error bars represent the standard deviation. P-values were calculated using unpaired, two-tailed t tests comparing each mutant to wild type (n.s., not significant; ^∗∗^, P ≤ 0.01; ^∗∗∗∗^, P ≤ 0.0001). For full statistical reporting, refer to Table 4. (C) Left: Representative thin sections of WT and *cco* mutant biofilms, stained with lectin and imaged by fluorescence microscopy. Biofilms were grown for two days before sampling. Right: Relative quantification of lectin stain signal intensity. Coloration of strain names in the left panel provides a key for the plotted data, and the y-axis in the right panel provides a scale bar for the left panel. Lectin-staining images and values are representative of four biological replicates.

We hypothesized that one or more of the CcoN subunits encoded by the PA14 genome is required for phenazine reduction and tested this by measuring the redox potential over depth for a series of *cco* mutants (**Figure 5B, top**). We saw very little reduction of phenazines in the Δ*cco1cco2* colony, suggesting that *cbb*_3_-type oxidases are required for this activity. In contrast, the mutant lacking the catalytic subunits of Cco1 and Cco2, Δ*N1*Δ*N2*, showed a redox profile similar to the wild type, indicating that isoforms containing one or both of the orphan CcoN subunits could support phenazine reduction activity. Indeed, although redox profiles obtained for the Δ*N1*Δ*N2* and Δ*N4* mutants were similar to those obtained for the wild type, the redox profile of the Δ*N1*Δ*N2*Δ*N4* mutant recapitulated that of Δ*cco1cco2.* These results indicate redundancy in the roles of some of the CcoN subunits. Consistent with this, Δ*N1*Δ*N4* showed an intermediate defect in phenazine reduction. Extraction and measurement of phenazines released from wild-type and *cco* mutant biofilms showed that variations in redox profiles could not be attributed to differences in phenazine production (**Figure 5— figure supplement 1**).

Our group has previously shown that a Δ*phz* mutant compensates for its lack of phenazines by forming thinner colonies, thus limiting the development of the hypoxic subzone seen in the wild type (Dietrich et al. 2013). We therefore hypothesized that mutants unable to reduce phenazines would likewise result in thinner colonies. Indeed, we observed that the *cco* mutants that lacked phenazine reduction profiles in the top panel of **Figure 5B** produced biofilms that were significantly thinner than wild-type and comparable to that of the Δ*phz* mutant (**Figure 5B, bottom**).

### Wild-type and cco mutant colony biofilms show increased matrix production at a consistent depth

We have recently demonstrated that extracellular matrix production, a hallmark of biofilm formation, is regulated by redox state in PA14 colony biofilms. Increased matrix production correlates with the accumulation of reducing power (as indicated by higher cellular NADH/NAD^+^ ratios) due to electron acceptor limitation and is visible in the hypoxic region of Δ*phz* colonies (Dietrich et al. 2013; Okegbe et al. 2017). The morphologies of our *cco* mutants (**Figure 2A**) suggest that matrix production can also be induced by respiratory chain dysfunction, which may be linked to defects in phenazine utilization (**Figure 5B**). To further examine the relationships between Cco isoforms and redox imbalance in biofilms, we prepared thin sections from two day-old colonies and stained with fluorescein-labeled lectin, which binds preferentially to the Pel polysaccharide component of the matrix (Jennings et al. 2015). Consistent with their similar gross morphologies, the wild-type and Δ*N1*Δ*N2* biofilms showed similar patterns of staining, with a faint band of higher intensity at a depth of ~40 μm (**Figure 5C**). Δ*N4* also showed a similar pattern, with a slightly higher intensity of staining in this band. Δ*N1*Δ*N2*Δ*N4* and Δ*cco1cco2* showed more staining throughout each sample, with wider bands of greater intensity at the ~40 μm point. These data suggest that deletion of the Cco complexes leads to a more reduced biofilm, which induces production of more matrix, and that CcoN4 contributes significantly to maintaining redox homeostasis when O_2_ is limiting.

### ccoN4 contributes to P. aeruginosa virulence in a C. elegans slow killing model

We have previously shown that a mutant defective in biofilm-specific phenazine production, which also shows altered colony morphology (Dietrich et al. 2008; Dietrich et al. 2013), exhibits decreased virulence (Recinos et al. 2012). We and others have suggested that one way in which phenazines could contribute to virulence is by acting as electron acceptors to balance the intracellular redox state in the hypoxic conditions that are encountered during infection (Price-Whelan et al. 2006; Newman 2008; Dietrich et al. 2013). Because CcoN4 is required for wildtype biofilm architecture and respiration (**Figures 2A, 2C, and 5C**), we hypothesized that it could also contribute to virulence. To test this, we conducted virulence assays using the nematode *Caenorhabditis elegans* as a host. It has been shown that *P. aeruginosa* is pathogenic to *C. elegans* and that the slow killing assay mimics an infection-like killing of *C. elegans* by the bacterium (Tan et al. 1999). While Δ*N1*Δ*N2* killed with wild type-like kinetics, Δ*N1*Δ*N2*Δ*N4* and Δ*cco1cco2* both showed comparably-impaired killing relative to wild-type PA14 (**Figure 6**).

**Figure 6.**
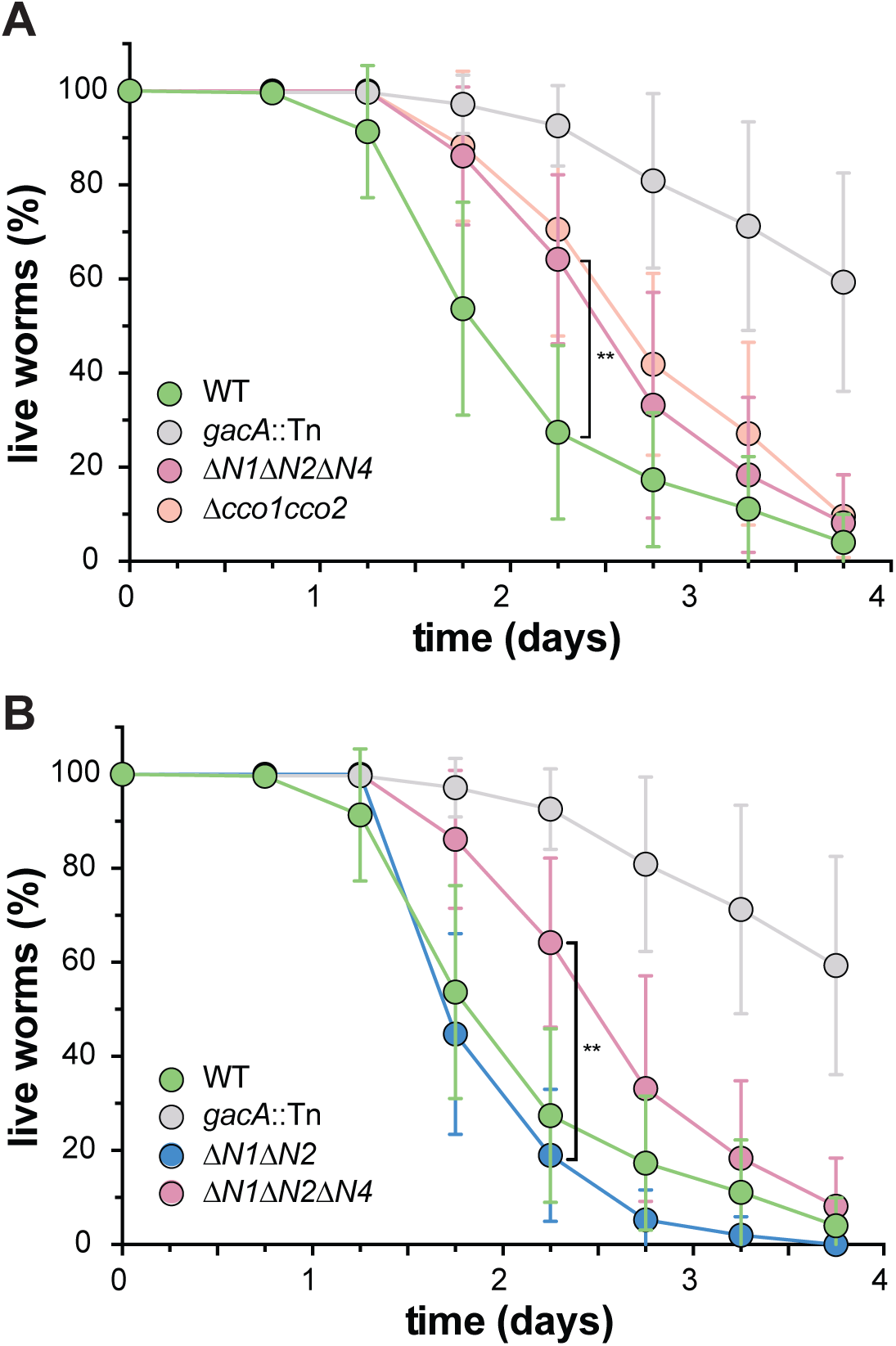
CcoN4-containing isoform(s) make unique contributions to PA14 virulence. Slow-killing kinetics of WT, *gacA*, and various *cco* mutant strains in the nematode *Caenorhabditis elegans.* Nearly 100% of the *C. elegans* population exposed to WT PA14 is killed after four days of exposure to the bacterium, while a mutant lacking GacA, a regulator that controls expression of virulence genes in *P aeruginosa*, shows decreased killing, with ~50% of worms alive four days post-exposure. (A) Δ*N1*Δ*N2*Δ*N4* and Δ*cco1cco2* show comparably attenuated pathogenicity relative to WT. Error bars represent the standard deviation of at least six biological replicates. At 2.25 days post-exposure, significantly less *C. elegans* were killed by Δ*N1*Δ*N2*Δ*N4* than by WT (unpaired two-tailed t test; p = 0.0022). (B) Δ*N1*Δ*N2* displays only slightly reduced pathogenicity when compared to WT. At 2.25 days post-exposure, significantly more *C. elegans* were killed by Δ*N1*Δ*N2* than by Δ*N1*Δ*N2*Δ*N4* (unpaired two-tailed t test; p = 0.003). For full statistical reporting, refer to Table 4. Error bars represent the standard deviation of at least four biological replicates, each with a starting sample size of 30-35 worms per replicate.

## DISCUSSION

Biofilm formation contributes to *P. aeruginosa* pathogenicity and persistence during different types of infections, including the chronic lung colonizations seen in individuals with cystic fibrosis (Tolker-Nielsen 2014; Rybtke et al. 2015). The conditions found within biofilm microenvironments are distinct from those in well-mixed liquid cultures with respect to availability of electron donors and acceptors. We have previously described the roles of phenazines, electron-shuttling antibiotics produced by *P. aeruginosa*, in biofilm-specific metabolism. In this study, we focused on *P. aeruginosa’s* large complement of genes encoding *cbb*3-type cytochrome oxidase subunits and set out to test their contributions to metabolic electron flow in biofilms.

The *P. aeruginosa* genome contains four different homologs of *ccoN*, encoding the catalytic subunit of *cbb*_3_-type oxidase. Only two of these (*ccoN1* and *ccoN2*) are co-transcribed with a *ccoO* homolog, encoding the other critical component of an active *cbb*_3_-type oxidase (**Figure 1B**). However, genetic studies have demonstrated that all four versions of CcoN can form functional complexes when expressed with either of the two CcoO homologs (Hirai et al. 2016). In well-mixed liquid cultures, mutants lacking the “orphan” subunits do not show growth defects (**Figure 2C**) (Hirai et al. 2016). We were therefore surprised to find that the Δ*N4* mutant showed a unique morphotype in a colony biofilm assay (**Figure 2A**). We have applied this assay extensively in our studies of the mechanisms underlying cellular redox balancing and sensing and noted that the phenotype of Δ*N4* was similar to that of mutants with defects in electron shuttling and redox signaling (Dietrich et al. 2013; Okegbe et al. 2017).

We characterized the effects of a Δ*N4* mutation on biofilm physiology through a series of assays. In well-mixed liquid cultures, Δ*colcco2* shows a growth phenotype similar to that of Δ*N1*Δ*N2*, suggesting that the subunits of the Cco1 and Cco2 oxidases do not form heterocomplexes with CcoN4 or that such complexes do not contribute to growth under these conditions. Consistent with this, deleting *ccoN4* in the Δ*N1*Δ*N2* background has no effect on growth. However, in biofilm-based experiments, we found that deleting *N4* alone was sufficient to cause an altered morphology phenotype (**Figure 2A**), and that deleting *N4* in either a Δ*N1* or a Δ*N1*Δ*N2* background profoundly affected biofilm physiology. These experiments included quantification of respiratory activity in colonies, in which deletion of CcoN4 led to a significant decrease (**Figure 2B**); biofilm co-culturing, in which CcoN4 was required for competitive fitness (**Figure 3A and B**); redox profiling, which showed that CcoN4 can contribute to phenazine reduction (**Figure 5B, top**); colony thickness measurements, which showed that CcoN4 is required for the formation of the hypoxic and anoxic zones (**Figure 5B, bottom**); and matrix profiling, which showed that CcoN4 contributes to the repression of Pel polysaccharide production (**Figure 5C**). The overlap in zones of expression between *cco1*, *cco2*, and *ccoN4Q4* seen in colony thin sections (**Figure 4**) implies that CcoN4 could form heterocomplexes with Cco1 and Cco2 subunits that span the depth of the colony and function to influence the physiology of *P. aeruginosa* biofilms in these ways.

Our results suggest that CcoN4 supports O_2_ and/or phenazine reduction specifically in biofilms. Using a strain engineered to produce GFP under control of the promoter for *ccoN4Q4*, we found that this locus is expressed throughout the biofilm depth, suggesting that CcoN4-containing isoforms could contribute to cytochrome *c* oxidation in both oxic and hypoxic zones (**Figure 4**). This constitutes a deviation from the previously published observation that these genes are specifically induced in hypoxic liquid cultures when compared to well-aerated ones (Alvarez-Ortega & Harwood 2007). We conclude that *ccoN4Q4* is uniquely induced by the conditions in the upper portion of the biofilm, where O_2_ is available as an electron acceptor. This regulation may contribute to the biofilm-specific role of the CcoN4 subunit.

The results described here can inform our model of how cells survive under distinct conditions in the microenvironments within biofilms. Previous work has shown that pyruvate fermentation can support survival of *P. aeruginosa* under anoxic conditions (Eschbach et al. 2004) and that phenazines facilitate this process (Glasser et al. 2014). Additional research suggests that phenazine reduction is catalyzed adventitiously by *P. aeruginosa* flavoproteins and dehydrogenases (Glasser et al. 2017). Our observation that *cbb*_3_-type cytochrome oxidases, particularly those containing the CcoN1 or CcoN4 subunits, are required for phenazine reduction in hypoxic biofilm subzones (**Figure 5B**) implicates the electron transport chain in utilization of these compounds. Various mechanisms of phenazine reduction and phenazine-related metabolisms may be relevant at different biofilm depths or depending on electron donor availability. Our results suggest that, in the colony biofilm system, enzyme complexes traditionally considered to be specific to oxygen reduction may contribute to anaerobic survival.

Because biofilm formation is often associated with colonization of and persistence in hosts, we tested whether *ccoN4* contributes to *P. aeruginosa* pathogenicity in *C. elegans.* Similar to our observations in biofilm assays, we found that the Δ*cco1cco2* mutant displayed a more severe phenotype than the Δ*N1*Δ*N2* mutant, suggesting that an orphan subunit can substitute for those encoded by the *cco1* and *cco2* operons. We also found that deleting *ccoN4* in Δ*N1*Δ*N2* led to a Δ*cco1cco2*-like phenotype, suggesting that CcoN4 is the subunit that can play this role (**Figure 6**). In host microenvironments where O_2_ is available, CcoN4-containing isoforms could contribute to its reduction. Additionally, in hypoxic zones, CcoN4-containing isoforms could facilitate the reduction of phenazines, enabling cellular redox balancing. Both of these functions would contribute to persistence of the bacterium within the host. The contributions of the Cco oxidases to *P. aeruginosa* pathogenicity raise the possibility that compounds interfering with Cco enzyme function could be effective therapies for these infections. Such drugs would be attractive candidates due to their specificity for bacterial respiratory chains and, as such, would not affect the host’s endogenous respiratory enzymes.

Our discovery that an orphan *cbb*_3_-type oxidase subunit contributes to growth in biofilms further expands the picture of *P. aeruginosa’s* remarkable respiratory flexibility. Beyond modularity at the level of the terminal enzyme complex (e.g., utilization of an *aa*_3_- vs. a *cbb*_3_-type oxidase), the activity of *P. aeruginosa’s* respiratory chain is further influenced by substitution of orphan *cbb*_3_-type catalytic subunits for native ones. Utilization of CcoN4-containing isoforms promotes phenazine reduction activity and may influence aerobic respiration in *P. aeruginosa* biofilms. For the exceptional species that contain orphan *cbb*_3_-type catalytic subunits, this fine level of control could be particularly advantageous during growth and survival in environments covering a wide range of electron acceptor availability (Cowley et al. 2015).

## MATERIALS AND METHODS

### Bacterial strains and growth conditions

*P. aeruginosa* strain UCBPP-PA14 (Rahme et al. 1995) was routinely grown in lysogeny broth (LB; 1% tryptone, 1% NaCl, 0.5% yeast extract) (Bertani 2004) at 37°C with shaking at 250 rpm unless otherwise indicated. Overnight cultures were grown for 12-16 hours. For genetic manipulation, strains were typically grown on LB solidified with 1.5% agar. Strains used in this study are listed in Table 3. In general, liquid precultures served as inocula for experiments. Overnight precultures for biological replicates were started from separate clonal source colonies on streaked agar plates. For technical replicates, a single preculture served as the source inoculum for subcultures.

### Construction of mutant *P. aeruginosa* strains

For making markerless deletion mutants in *P. aeruginosa* PA14 (Table 3) 1 kb of flanking sequence from each side of the target gene were amplified using the primers listed in Table 1 and inserted into pMQ30 through gap repair cloning in *Saccharomyces cerevisiae* InvSc1 (Shanks et al. 2006). Each plasmid listed in Table 2 was transformed into *Escherichia coli* strain UQ950, verified by restriction digests, and moved into PA14 using biparental conjugation. PA14 single recombinants were selected on LB agar plates containing 100 μg/ml gentamicin. Double recombinants (markerless deletions) were selected on LB without NaCl and modified to contain 10% sucrose. Genotypes of deletion mutants were confirmed by PCR. Combinatorial mutants were constructed by using single mutants as hosts for biparental conjugation, with the exception of Δ*cco1cco2*, which was constructed by deleting the *cco1* and *cco2* operons simultaneously as one fragment. *ccoN4* complementation strains were made in the same manner, using primers LD438 and LD441 listed in Table 1 to amplify the coding sequence of *ccoN4*, which was verified by sequencing and complemented back into the site of the deletion.

**Table 1.**
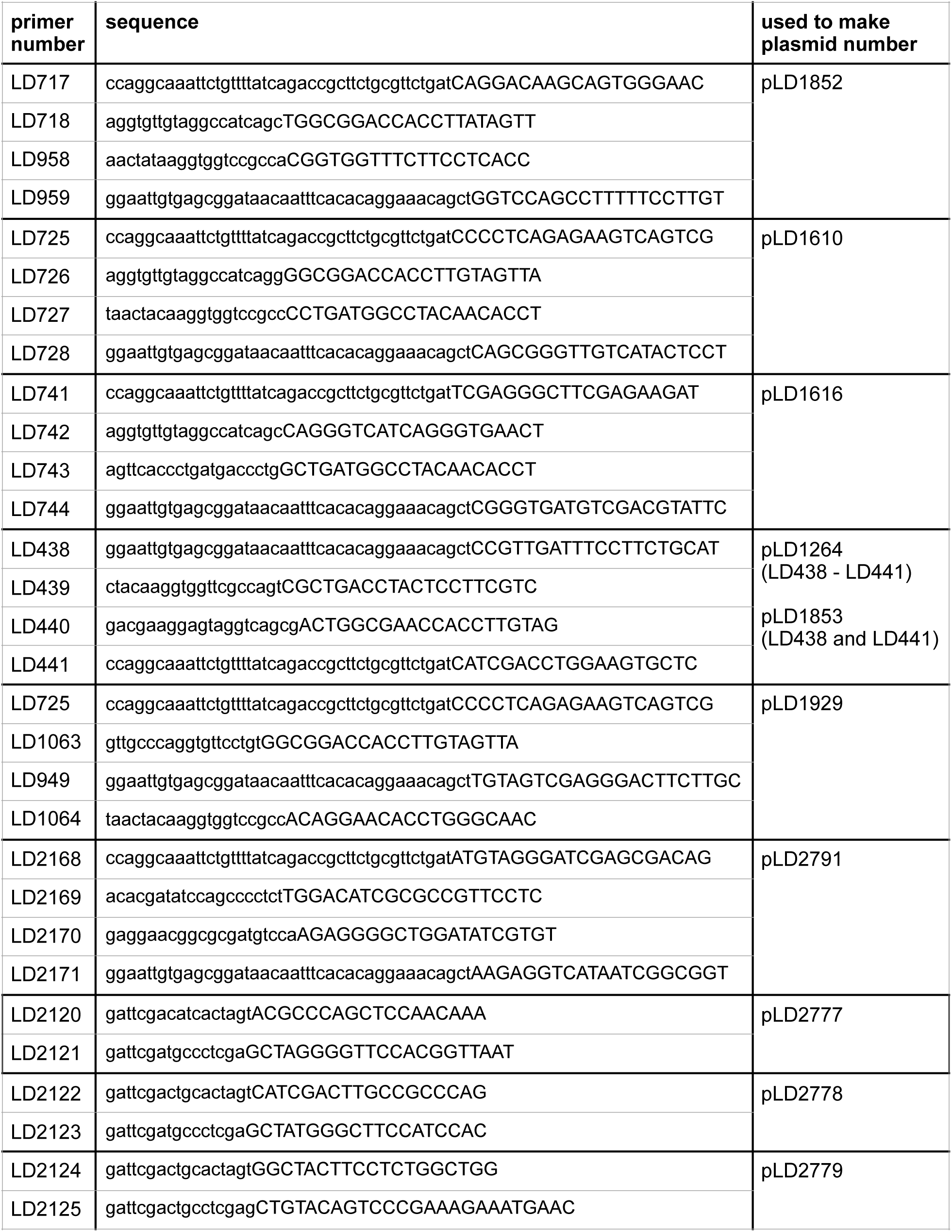
Primers used in this study.

**Table 2.** Plasmids used in this study.

| plasmid | description | source |
| --- | --- | --- |
| pMQ30 | 7.5 kb mobilizable vector; oriT, sacB, Gm<sup>R</sup>. | Shanks et al. 2006 |
| pAKN69 | Contains mini-Tn7(Gm)PA1/04/03::eyfp fusion. | Lambertsen et al. 2004 |
| pLD2722 | Gm<sup>R</sup>, Tet<sup>R</sup> flanked by Flp recombinase target (FRT) sites to resolve out resistance cassettes. | this study |
| pFLP2 | Site-specific excision vector with cl857-controlled FLP recombinase encoding sequence, sacB, Ap<sup>R</sup>. | Hoang et al. 1998 |
| pLD1852 | ΔccoN1 PCR fragment introduced into pMQ30 by gap repair cloning in yeast strain InvSc1. | this study |
| pLD1610 | ΔccoN2 PCR fragment introduced into pMQ30 by gap repair cloning in yeast strain InvSc1. | this study |
| pLD1616 | ΔccoN3 PCR fragment introduced into pMQ30 by gap repair cloning in yeast strain InvSc1. | this study |
| pLD1264 | ΔccoN4 PCR fragment introduced into pMQ30 by gap repair cloning in yeast strain InvSc1. | this study |
| pLD1929 | Δcco1 cco2 PCR fragment introduced into pMQ30 by gap repair cloning in yeast strain InvSc1. | this study |
| pLD2791 | Δhcn PCR fragment introduced into pMQ30 by gap repair cloning in yeast strain InvSc1. | this study |
| pLD1853 | Full genomic sequence of ccoN4 PCR fragment introduced into pMQ30 by gap repair cloning in yeast strain InvSc1. Verified by sequencing. | this study |
| pLD2777 | PCR-amplified cco1 promoter ligated into pSEK103 using SpeI and XhoI. | this study |
| pLD2778 | PCR-amplified cco2 promoter ligated into pSEK103 using SpeI and XhoI. | this study |
| pLD2779 | PCR-amplified ccoN4 promoter ligated into pSEK103 using SpeI and XhoI. | this study |

**Table 3.**
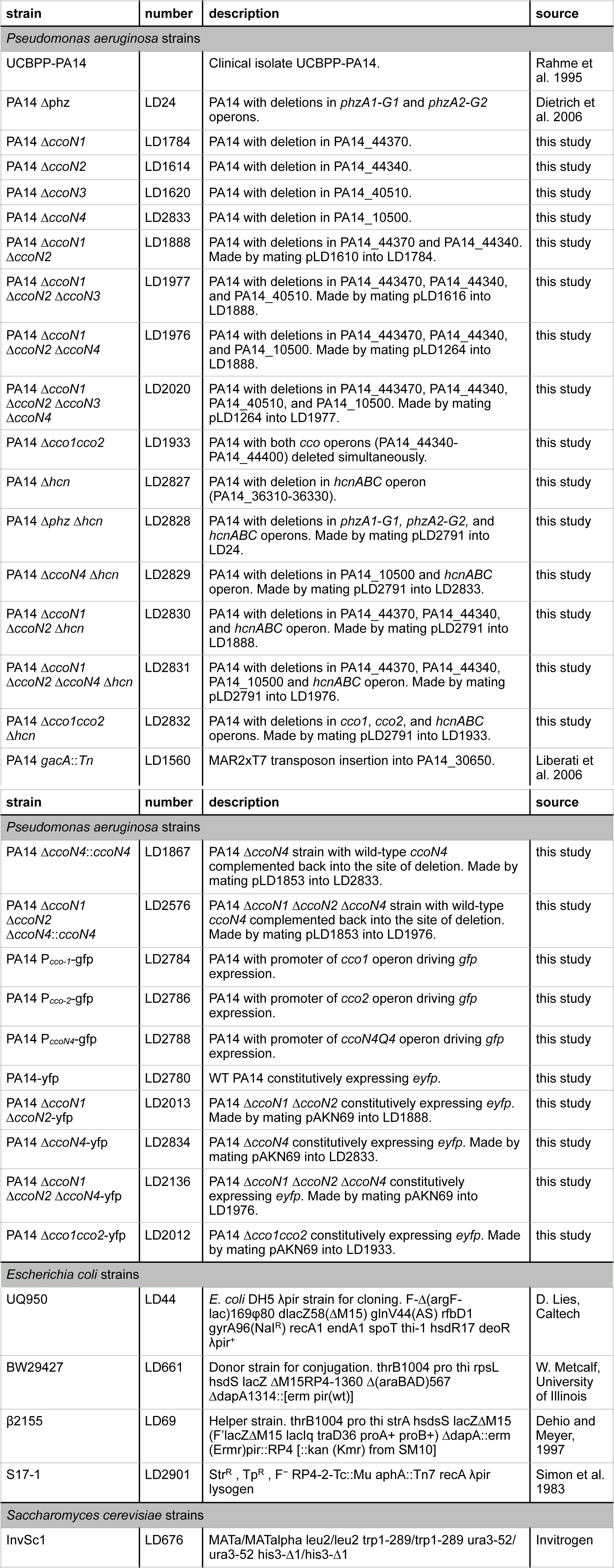
Strains used in this study.

| strain | number | description | source |
| --- | --- | --- | --- |
| *Pseudomonas aeruginosa* strains |  |  |  |
| UCBPP-PA14 |  | Clinical isolate UCBPP-PA14. | Rahme et al. 1995 |
| PA14 Δphz | LD24 | PA14 with deletions in *phzA1-G1* and *phzA2-G2* operons. | Dietrich et al. 2006 |
| PA14 ΔccoN1 | LD1784 | PA14 with deletion in PA14_44370. | this study |
| PA14 ΔccoN2 | LD1614 | PA14 with deletion in PA14_44340. | this study |
| PA14 ΔccoN3 | LD1620 | PA14 with deletion in PA14_40510. | this study |
| PA14 ΔccoN4 | LD2833 | PA14 with deletion in PA14_10500. | this study |
| PA14 ΔccoN1 ΔccoN2 | LD1888 | PA14 with deletions in PA14_44370 and PA14_44340. Made by mating pLD1610 into LD1784. | this study |
| PA14 ΔccoN1 ΔccoN2 ΔccoN3 | LD1977 | PA14 with deletions in PA14_443470, PA14_44340, and PA14_40510. Made by mating pLD1616 into LD1888. | this study |
| PA14 ΔccoN1 ΔccoN2 ΔccoN4 | LD1976 | PA14 with deletions in PA14_443470, PA14_44340, and PA14_10500. Made by mating pLD1264 into LD1888. | this study |
| PA14 ΔccoN1 ΔccoN2 ΔccoN3 ΔccoN4 | LD2020 | PA14 with deletions in PA14_443470, PA14_44340, PA14_40510, and PA14_10500. Made by mating pLD1264 into LD1977. | this study |
| PA14 Δcco1cco2 | LD1933 | PA14 with both *cco* operons (PA14_44340-PA14_44400) deleted simultaneously. | this study |
| PA14 Δhcn | LD2827 | PA14 with deletion in *hcnABC* operon (PA14_36310-36330). | this study |
| PA14 Δphz Δhcn | LD2828 | PA14 with deletions in *phzA1-G1*, *phzA2-G2*, and *hcnABC* operons. Made by mating pLD2791 into LD24. | this study |
| PA14 ΔccoN4 Δhcn | LD2829 | PA14 with deletions in PA14_10500 and *hcnABC* operon. Made by mating pLD2791 into LD2833. | this study |
| PA14 ΔccoN1 ΔccoN2 Δhcn | LD2830 | PA14 with deletions in PA14_44370, PA14_44340, and *hcnABC* operon. Made by mating pLD2791 into LD1888. | this study |
| PA14 ΔccoN1 ΔccoN2 ΔccoN4 Δhcn | LD2831 | PA14 with deletions in PA14_44370, PA14_44340, PA14_10500 and *hcnABC* operon. Made by mating pLD2791 into LD1976. | this study |
| PA14 Δcco1cco2 Δhcn | LD2832 | PA14 with deletions in *cco1*, *cco2*, and *hcnABC* operons. Made by mating pLD2791 into LD1933. | this study |
| PA14 *gacA::Tn* | LD1560 | MAR2xT7 transposon insertion into PA14_30650. | Liberati et al. 2006 |

**Table 3 (continued). Strains used in this study.**
| strain | number | description | source |
| --- | --- | --- | --- |
| *Pseudomonas aeruginosa* strains |  |  |  |
| PA14 $\Delta ccoN4::ccoN4$ | LD1867 | PA14 $\Delta ccoN4$ strain with wild-type *ccoN4* complemented back into the site of deletion. Made by mating pLD1853 into LD2833. | this study |
| PA14 $\Delta ccoN1$<br>$\Delta ccoN2$<br>$\Delta ccoN4::ccoN4$ | LD2576 | PA14 $\Delta ccoN1$ $\Delta ccoN2$ $\Delta ccoN4$ strain with wild-type *ccoN4* complemented back into the site of deletion. Made by mating pLD1853 into LD1976. | this study |
| PA14 P<sub>cco-1</sub>-gfp | LD2784 | PA14 with promoter of *cco1* operon driving *gfp* expression. | this study |
| PA14 P<sub>cco-2</sub>-gfp | LD2786 | PA14 with promoter of *cco2* operon driving *gfp* expression. | this study |
| PA14 P<sub>ccoN4Q4</sub>-gfp | LD2788 | PA14 with promoter of *ccoN4Q4* operon driving *gfp* expression. | this study |
| PA14-yfp | LD2780 | WT PA14 constitutively expressing *eyfp*. | this study |
| PA14 $\Delta ccoN1$<br>$\Delta ccoN2$-yfp | LD2013 | PA14 $\Delta ccoN1$ $\Delta ccoN2$ constitutively expressing *eyfp*. Made by mating pAKN69 into LD1888. | this study |
| PA14 $\Delta ccoN4$-yfp | LD2834 | PA14 $\Delta ccoN4$ constitutively expressing *eyfp*. Made by mating pAKN69 into LD2833. | this study |
| PA14 $\Delta ccoN1$<br>$\Delta ccoN2$ $\Delta ccoN4$-yfp | LD2136 | PA14 $\Delta ccoN1$ $\Delta ccoN2$ $\Delta ccoN4$ constitutively expressing *eyfp*. Made by mating pAKN69 into LD1976. | this study |
| PA14 $\Delta cco1cco2$-yfp | LD2012 | PA14 $\Delta cco1cco2$ constitutively expressing *eyfp*. Made by mating pAKN69 into LD1933. | this study |
| *Escherichia coli* strains |  |  |  |
| UQ950 | LD44 | *E. coli* DH5 λpir strain for cloning. F-Δ(argF-lac)169ϕ80 dlacZ58(ΔM15) glnV44(AS) rfbD1 gyrA96(Nal<sup>R</sup>) recA1 endA1 spoT thi-1 hsdR17 deoR λpir<sup>+</sup> | D. Lies, Caltech |
| BW29427 | LD661 | Donor strain for conjugation. thrB1004 pro thi rpsL hsdS lacZ ΔM15RP4-1360 $\Delta$(araBAD)567 ΔdapA1314::[erm pir(wt)] | W. Metcalf, University of Illinois |
| β2155 | LD69 | Helper strain. thrB1004 pro thi strA hsdS lacZΔM15 (F'*lacZ*ΔM15 *lacIq* traD36 proA<sup>+</sup> proB<sup>+</sup>) ΔdapA::erm (Erm<sup>r</sup>)pir::RP4 [::kan (Kmr) from SM10] | Dehio and Meyer, 1997 |
| S17-1 | LD2901 | Str<sup>R</sup>, Tp<sup>R</sup>, F<sup>-</sup> RP4-2-Tc::Mu aphA::Tn7 recA λpir lysogen | Simon et al. 1983 |
| *Saccharomyces cerevisiae* strains |  |  |  |
| InvSc1 | LD676 | MATa/MATalpha leu2/leu2 trp1-289/trp1-289 ura3-52/ura3-52 his3-Δ1/his3-Δ1 | Invitrogen |

**Table 4.** Statistical analysis.

| Figure 2B | number of values (biological replicates) | mean | median | SD | SEM | lower 95% confidence interval of mean | upper 95% confidence interval of mean |
| --- | --- | --- | --- | --- | --- | --- | --- |
| WT | 5 | 73.22 | 72.94 | 3.387 | 1.515 | 69.02 | 77.43 |
| $\Delta N4$ | 5 | 68.97 | 70.6 | 6.44 | 2.88 | 60.97 | 76.96 |
| $\Delta N1\Delta N2$ | 5 | 52.18 | 50.46 | 5.142 | 2.3 | 45.79 | 58.56 |
| $\Delta N1\Delta N4$ | 5 | 11.57 | 12.42 | 2.011 | 0.8991 | 9.074 | 14.07 |
| $\Delta N1\Delta N2\Delta N4$ | 5 | 0.001958 | 0.001117 | 0.001696 | 0.0007586 | -0.0001481 | 0.004064 |
| $\Delta cco1cco2$ | 5 | 0.001367 | 0.0008644 | 0.001237 | 0.0005532 | -0.0001686 | 0.002903 |
| **t-test** | **P value** | **P value summary** |  |  |  |  |  |
| WT vs. $\Delta N4$ | 0.2273 | ns | | | | | |
| WT vs. $\Delta N1\Delta N2$ | <0.0001 | **** | | | | | |
| WT vs. $\Delta N1\Delta N4$ | <0.0001 | **** | | | | | |
| WT vs. $\Delta N1\Delta N2\Delta N4$ | <0.0001 | **** | | | | | |
| WT vs. $\Delta cco1cco2$ | <0.0001 | **** | | | | | |

| Figure 3A | number of values (biological replicates) | mean | median | SD | SEM | lower 95% confidence interval of mean | upper 95% confidence interval of mean |
| --- | --- | --- | --- | --- | --- | --- | --- |
| WT-YFP | 12 | 54.95 | 54.92 | 4.387 | 1.266 | 52.16 | 57.74 |
| $\Delta N4$-YFP | 3 | 29.92 | 30.83 | 2.234 | 1.29 | 24.37 | 35.46 |
| $\Delta N1\Delta N2$-YFP | 3 | 30.49 | 31.91 | 3.527 | 2.036 | 21.73 | 39.25 |
| $\Delta N1\Delta N2\Delta N4$-YFP | 3 | 4.408 | 4.296 | 3.23 | 1.865 | -3.617 | 12.43 |
| $\Delta cco1cco2$-YFP | 3 | 7.097 | 5.306 | 4.093 | 2.363 | -3.072 | 17.27 |
| **t-test** | **P value** | **P value summary** |  |  |  |  |  |
| WT-YFP vs. $\Delta N4$-YFP | <0.0001 | **** | | | | | |
| WT-YFP vs. $\Delta N1\Delta N2$-YFP | <0.0001 | **** | | | | | |
| $\Delta N1\Delta N2$-YFP vs. $\Delta N1\Delta N2\Delta N4$-YFP | 0.0007 | *** | | | | | |
| $\Delta N1\Delta N2$-YFP vs. $\Delta cco1cco2$-YFP | 0.0017 | *** | | | | | |

**Table 4 (continued). Statistical analysis.**
| Figure 3– figure supplement 1A | number of values (biological replicates) | mean | median | SD | SEM | lower 95% confidence interval of mean | upper 95% confidence interval of mean |
| --- | --- | --- | --- | --- | --- | --- | --- |
| WT | 12 | 45.05 | 45.08 | 4.387 | 1.266 | 42.26 | 47.84 |
| $\Delta N4$ | 3 | 28.22 | 31.31 | 7.442 | 4.297 | 9.731 | 46.71 |
| $\Delta N1\Delta N2$ | 3 | 27.81 | 28.57 | 2.514 | 1.451 | 21.56 | 34.05 |
| $\Delta N1\Delta N2\Delta N4$ | 3 | 7.002 | 6.973 | 0.7508 | 0.4335 | 5.137 | 8.867 |
| $\Delta cco1cco2$ | 3 | 5.38 | 4.183 | 2.146 | 1.239 | 0.05034 | 10.71 |
| t-test | P value | P value summary |  |  |  |  |  |
| WT vs. $\Delta N4$ | 0.0002 | *** | | | | | |
| WT vs. $\Delta N1\Delta N2$ | <0.0001 | **** | | | | | |
| $\Delta N1\Delta N2$ vs. $\Delta N1\Delta N2\Delta N4$ | 0.0002 | *** | | | | | |
| $\Delta N1\Delta N2$ vs. $\Delta cco1cco2$ | 0.0003 | *** | | | | | |

| Figure 5 | number of values (biological replicates) | mean | median | SD | SEM | lower 95% confidence interval of mean | upper 95% confidence interval of mean |
| --- | --- | --- | --- | --- | --- | --- | --- |
| WT | 8 | 150.3 | 151.2 | 10.31 | 3.644 | 141.7 | 158.9 |
| $\Delta N1\Delta N2$ | 4 | 139.3 | 137.6 | 12.33 | 6.166 | 119.6 | 158.9 |
| $\Delta N4$ | 7 | 131.9 | 127.8 | 8.915 | 3.369 | 123.7 | 140.2 |
| $\Delta N1\Delta N4$ | 4 | 99.96 | 99.34 | 2.726 | 1.363 | 95.62 | 104.3 |
| $\Delta cco1cco2$ | 4 | 95.19 | 95.56 | 1.559 | 0.7793 | 92.71 | 97.67 |
| $\Delta N1\Delta N2\Delta N4$ | 4 | 102.8 | 99.79 | 8.664 | 4.332 | 88.98 | 116.6 |
| $\Delta phz$ | 7 | 84.98 | 84.23 | 10.93 | 4.131 | 74.87 | 95.09 |
| t-test | P value | P value summary |  |  |  |  |  |
| WT vs. $\Delta N1\Delta N2$ | 0.1302 | ns | | | | | |
| WT vs. $\Delta N4$ | 0.0028 | ** | | | | | |
| WT vs. $\Delta N1\Delta N4$ | <0.0001 | **** | | | | | |
| WT vs. $\Delta cco1cco2$ | <0.0001 | **** | | | | | |
| WT vs. $\Delta N1\Delta N2\Delta N4$ | <0.0001 | **** | | | | | |
| WT vs. $\Delta phz$ | <0.0001 | **** | | | | | |

**Table 4 (continued). Statistical analysis.-1**
| **Figure 6** | **number of values**<br>**(biological replicates)** | **mean** | **median** | **SD** | **SEM** | **lower 95% confidence interval of mean** | **upper 95% confidence interval of mean** |
| --- | --- | --- | --- | --- | --- | --- | --- |
| **WT** | 9 | 27.44 | 39 | 18.48 | 6.16 | 13.24 | 41.65 |
| ***gacA::Tn*** | 9 | 92.56 | 93 | 8.546 | 2.849 | 85.99 | 99.12 |
| **$\Delta N1\Delta N2$** | 4 | 19 | 21.5 | 14.07 | 7.036 | -3.39 | 41.39 |
| **$\Delta N1\Delta N2\Delta N4$** | 6 | 64.17 | 68 | 18 | 7.35 | 45.27 | 83.06 |
| **$\Delta cco1cco2$** | 9 | 70.56 | 76 | 22.69 | 7.565 | 53.11 | 88 |
| **t-test** | **P value** | **P value summary** |  |  |  |  |  |
| **WT vs. $\Delta N1\Delta N2\Delta N4$** | 0.0022 | ** |  |  |  |  |  |
| **$\Delta N1\Delta N2$ vs. $\Delta N1\Delta N2\Delta N4$** | 0.0030 | ** |  |  |  |  |  |
| **WT vs. $\Delta N1\Delta N2$** | 0.4362 | ns |  |  |  |  |  |

### Colony biofilm morphology assays

Overnight precultures were diluted 1:100 in LB (Δ*N1*Δ*N2*, Δ*N1*Δ*N2*Δ*N3*, Δ*N1*Δ*N2*Δ*N4*, Δ*N1*Δ*N2*Δ*N4*Δ*N3*, Δ*N1*Δ*N2*Δ*N4::N4*, Δ*cco1cco2*, Δ*N1*Δ*N2*Δ*hcn*, Δ*N1*Δ*N2*Δ*N4*Δ*hcn*, and Δ*cco1cco2*Δ*hcn* were diluted 1:50) and grown to mid-exponential phase (OD at 500 nm ≈ 0.5). Ten microliters of subcultures were spotted onto 60 mL of colony morphology medium (1% tryptone, 1% agar containing 40 μg/ml Congo red dye and 20 μg/ml Coomassie blue dye) in a 10 cm x 10 cm x 1.5 cm square Petri dish (LDP D210-16). Plates were incubated for up to five days at 25°C with > 90% humidity (Percival CU-22L) and imaged daily using a Keyence VHX-1000 digital microscope. Images shown are representative of at least ten biological replicates. 3D images of biofilms were taken on day 5 of development using a Keyence VR-3100 wide-area 3D measurement system. *hcn* deletion mutants were imaged using a flatbed scanner (Epson E11000XL-GA) and are representative of at least three biological replicates

### TTC reduction assay

One microliter of overnight cultures (five biological replicates), grown as described above, was spotted onto a 1% tryptone, 1.5% agar plate containing 0.001% (w/v) TTC (2,3,5-triphenyl-tetrazolium chloride [Sigma-Aldrich T8877]) and incubated in the dark at 25°C for 24 hours. Spots were imaged using a scanner (Epson E11000XL-GA) and TTC reduction, normalized to colony area, was quantified using Adobe Photoshop CS5. Colorless TTC undergoes an irreversible color change to red when reduced. Pixels in the red color range were quantified and normalized to colony area using Photoshop CS5.

### Liquid culture growth assays

**(i)** Overnight precultures were diluted 1:100 (Δ*N1*Δ*N2*Δ*N4* and Δ*cco1cco2* were diluted 1:50) in 1% tryptone in a polystyrene 96-well plate (Greiner Bio-One 655001) and grown for 2 hours (OD_500nm_ ≈ 0.2). These cultures were then diluted 100-fold in 1% tryptone in a clear, flat-bottom polystyrene 96-well plate (VWR 82050-716) and incubated at 37°C with continuous shaking on the medium setting in a Biotek Synergy 4 plate reader. Growth was assessed by taking OD_500nm_ readings every thirty minutes for at least 24 hours. **(ii) *hcn* mutants:** Overnight precultures were diluted 1:100 (Δ*N1*Δ*N2*Δ*N4*Δ*hcn* and Δ*cco1cco2*Δ*hcn* were diluted 1:50) in MOPS minimal medium (50 mM 4-morpholinepropanesulfonic acid (pH 7.2), 43 mM NaCl, 93 mM NH4Cl, 2.2 mM KH2PO4, 1 mM MgSO4•7H2O, 1 μg/ml FeSO4•7H2O, 20 mM sodium succinate hexahydrate) and grown for 2.5 hours until OD at 500 nm ≈ 0.1). These cultures were then diluted 100-fold in MOPS minimal medium in a clear, flat-bottom polystyrene 96-well plate and incubated at 37°C with continuous shaking on the medium setting in a Biotek Synergy 4 plate reader. Growth was assessed by taking OD readings at 500 nm every thirty minutes for at least 24 hours. **(iii) Terminal oxidase reporters:** Overnight precultures were grown in biological triplicate; each biological triplicate was grown in technical duplicate. Overnight precultures were diluted 1:100 in 1% tryptone and grown for 2.5 hours until OD at 500 nm ≈ 0.1. These cultures were then diluted 100-fold in 1% tryptone in a clear-bottom, polystyrene black 96-well plate (VWR 82050-756) and incubated at 37°C with continuous shaking on the medium setting in a Biotek Synergy 4 plate reader. Expression of GFP was assessed by taking fluorescence readings at excitation and emission wavelengths of 480 nm and 510 nm, respectively, every hour for 24 hours. Growth was assessed by taking OD readings at 500 nm every 30 minutes for 24 hours. Growth and RFU values for technical duplicates were averaged to obtain the respective values for each biological replicate. RFU values for a strain without a promoter inserted upstream of the *gfp* gene were considered background and subtracted from the fluorescence values of each reporter.

### Competition assays

Overnight precultures of fluorescent (YFP-expressing) and non-fluorescent strains were diluted 1:100 in LB (Δ*N1*Δ*N2*, Δ*N1*Δ*N2*Δ*N4* and Δ*cco1cco2* were diluted 1:50) and grown to mid-exponential phase (OD at 500 nm ≈ 0.5). Exact OD at 500 nm values were read in a Spectronic 20D+ spectrophotometer (Thermo Scientific) and cultures were adjusted to the same OD. Adjusted cultures were then mixed in a 1:1 ratio of fluorescent:non-fluorescent cells and ten μl of this mixture were spotted onto colony morphology plates and grown for three days as described above. At specified time points, biofilms were collected, suspended in one ml of 1% tryptone, and homogenized on the “high” setting in a bead mill homogenizer (Omni Bead Ruptor 12); day one colonies were homogenized for 35 seconds while days two and three colonies were homogenized for 99 seconds. Homogenized cells were serially diluted and 10^-6^, 10^-7^, and 10^-8^ dilutions were plated onto 1% tryptone plates and grown overnight at 37 °C. Fluorescent colony counts were determined by imaging plates with a Typhoon FLA7000 fluorescent scanner (GE Healthcare) and percentages of fluorescent vs. non-fluorescent colonies were determined.

### Construction of terminal oxidase reporters

Translational reporter constructs for the Cco1, Cco2, and CcoN4Q4 operons were constructed using primers listed in Table 1. Respective primers were used to amplify promoter regions (500 bp upstream of the operon of interest), adding an SpeI digest site to the 5’ end of the promoter and an XhoI digest site to the 3’ end of the promoter. Purified PCR products were digested and ligated into the multiple cloning site of the pLD2722 vector, upstream of the *gfp* sequence. Plasmids were transformed into *E. coli* strain UQ950, verified by sequencing, and moved into PA14 using biparental conjugation with *E. coli* strain S17-1. PA14 single recombinants were selected on M9 minimal medium agar plates (47.8 mM Na_2_HPO_4_•7H_2_O, 22 mM KH_2_PO_4_, 8.6 mM NaCl, 18.6 mM NH_4_Cl, 1 mM MgSO_4_, 0.1 mM CaCl_2_, 20 mM sodium citrate dihydrate, 1.5% agar) containing 100 μg/ml gentamicin. The plasmid backbone was resolved out of PA14 using Flp-FRT recombination by introduction of the pFLP2 plasmid (Hoang et al. 1998) and selected on M9 minimal medium agar plates containing 300 μg/ml carbenicillin and further on LB agar plates without NaCl and modified to contain 10% sucrose. The presence of *gfp* in the final clones was confirmed by PCR.

### Thin sectioning analyses

Two layers of 1% tryptone with 1% agar were poured to depths of 4.5 mm (bottom) and 1.5 mm (top). Overnight precultures were diluted 1:100 (Δ*N1*Δ*N4*, Δ*N1*Δ*N2*Δ*N4*, Δ*cco1cco2* were diluted 1:50) in LB and grown until early-mid exponential phase, for two hours. Then five to ten μL of subculture were spotted onto the top agar layer and colonies were incubated in the dark at 25°C with >90% humidity [Percival CU-22L]). Colonies were grown for up to three days. At specified time points to be prepared for thin sectioning, colonies were covered by a 1.5 mm-thick 1% agar layer. Colonies sandwiched between two 1.5mm agar layers were lifted from the bottom layer and soaked for four hours in 50 mM L-lysine in phosphate buffered saline (PBS) (pH 7.4) at 4°C, then fixed in 4% paraformaldehyde, 50 mM L-lysine, PBS (pH 7.4) for four hours at 4°C, then overnight at 37 °C. Fixed colonies were washed twice in PBS and dehydrated through a series of ethanol washes (25%, 50%, 70%, 95%, 3x 100% ethanol) for 60 minutes each. Colonies were cleared via three 60-minute incubations in Histoclear-II (National Diagnostics HS-202) and infiltrated with wax via two separate washes of 100% Paraplast Xtra paraffin wax (Electron Microscopy Sciences; Fisher Scientific 50-276-89) for two hours each at 55°C, then colonies were allowed to polymerize overnight at 4°C. Tissue processing was performed using an STP120 Tissue Processor (Thermo Fisher Scientific 813150). Trimmed blocks were sectioned in ten μm-thick sections perpendicular to the plane of the colony using an automatic microtome (Thermo Fisher Scientific 905200ER), floated onto water at 45°C, and collected onto slides. Slides were air-dried overnight, heat-fixed on a hotplate for one hour at 45°C, and rehydrated in the reverse order of processing. Rehydrated colonies were immediately mounted in TRIS-Buffered DAPI:Fluorogel (Electron Microscopy Sciences; Fisher Scientific 50-246-93) and overlaid with a coverslip. Differential interference contrast (DIC) and fluorescent confocal images were captured using an LSM700 confocal microscope (Zeiss). Each strain was prepared in this manner in at least biological triplicates.

### Colony thickness measurements

Colonies were prepared for thin sectioning as described above, but growth medium was supplemented with 40 μg/ml Congo Red dye (VWR AAAB24310-14) and 20 μg/ml Coomassie Blue dye (Omnipur; VWR EM-3300). Colony height measurements were obtained from confocal DIC images using Fiji image processing software (Schindelin et al. 2012).

### Lectin staining

Two-day-old colonies were prepared for thin sectioning as described above. Rehydrated colonies were post-stained in 100 μg/mL fluorescein-labeled *Wisteria floribunda* lectin (Vector Laboratories FL-1351) in PBS before being washed twice in PBS, mounted in TRIS-buffered DAPI and overlaid with a coverslip. Fluorescent confocal images were captured using an LSM700 confocal microscope (Zeiss).

### Redox profiling of biofilms

A 25 μm-tip redox microelectrode and external reference (Unisense RD-25 and REF-RM) were used to measure the extracellular redox state of day two (~ 48 h) biofilms (grown as for the **colony biofilm morphology assays**). The redox microelectrode measures the tendency of a sample to take up or release electrons relative to the reference electrode, which is immersed in the same medium as the one on which the sample is grown. The redox microelectrode was calibrated according to manufacturer’s instructions using a two-point calibration to 1% quinhydrone in pH 4 buffer and 1% quinhydrone in pH 7 buffer. Redox measurements were taken every five μm throughout the depth of the biofilm using a micromanipulator (Unisense MM33) with a measurement time of three seconds and a wait time between measurements of five seconds. Profiles were recorded using a multimeter (Unisense) and the SensorTrace Profiling software (Unisense).

### Oxygen profiling of biofilms

A 25 μm-tip oxygen microsensor (Unisense OX-25) was used to measure oxygen concentrations within biofilms during the first two days of development, grown as described above. For oxygen profiling on three-day-old colonies (Figure 4), biofilms were grown as for the thin sectioning analyses. To calibrate the oxygen microsensor, a two-point calibration was used. The oxygen microsensor was calibrated first to atmospheric oxygen using a calibration chamber (Unisense CAL300) containing water continuously bubbled with air. The microsensor was then calibrated to a “zero” point using an anoxic solution of water thoroughly bubbled with N_2_; to ensure complete removal of all oxygen, N_2_ was bubbled into the calibration chamber for a minimum of 30 minutes before calibrating the microsensor to the zero calibration point. Oxygen measurements were then taken throughout the depth of the biofilm using a measurement time of three seconds and a wait time between measurements of five seconds. For six-hour-old colonies, a step size of one μm was used to profile through the entire colony; for 12-hour and 24-hour colonies, two μm; for 48-hour colonies, five μm. A micromanipulator (Unisense MM33) was used to move the microsensor within the biofilm and profiles were recorded using a multimeter (Unisense) and the SensorTrace Profiling software (Unisense).

### Phenazine quantification

Overnight precultures were diluted 1:10 in LB and spotted onto a 25mm 0.2 μm filter disk (pore size: 0.2 μm; GE Healthcare 110606) placed into the center of one 35 x 10 mm round Petri dish (Falcon 351008). Colonies were grown for two days in the dark at 25°C with >90% humidity. After two days of growth, each colony (with filter disk) was lifted off its respective plate and weighed. Excreted phenazines were then extracted from the agar medium overnight in five mL of 100% methanol (in the dark, nutating at room temperature). Three hundred μl of this overnight phenazine/methanol extraction were then filtered through a 0.22 μm cellulose Spin-X column (Thermo Fisher Scientific 07-200-386) and 200 μl of the flow-through were loaded into an HPLC vial. Phenazines were quantified using high-performance liquid chromatography (Agilent 1100 HPLC System) as described previously (Dietrich, Price-Whelan, et al. 2006; Sakhtah et al. 2016).

### C. elegans pathogenicity (slow killing) assays

Slow killing assays were performed as described previously (Tan et al. 1999; Powell & Ausubel 2008). Briefly, ten μl of overnight PA14 cultures (grown as described above) were spotted onto slow killing agar plates (0.3% NaCl, 0.35% Bacto-Peptone, 1 mM CaCl_2_, 1 mM MgSO_4_, 5 μg/ml cholesterol, 25 mM KPO_4_, 50 μg/ml FUDR, 1.7% agar) and plates were incubated for 24 hours at 37 °C followed by 48 hours at room temperature (~23°C). Larval stage 4 (L4) nematodes were picked onto the PA14-seeded plates and live/dead worms were counted for up to four days. Each plate was considered a biological replicate and had a starting sample size of 30-35 worms.

### Statistical analysis

Data analysis was performed using GraphPad Prism version 7 (GraphPad Software, La Jolla California USA). Values are expressed as mean ± SD. Statistical significance of the data presented was assessed with the two-tailed unpaired Student’s t-test. Values of P ≤ 0.05 were considered significant (∗, P ≤ 0.05; ∗∗, P ≤ 0.01; ∗∗∗, P ≤0.001; ∗∗∗, P ≤ 0.0001 ∗∗∗∗, P ≤ 0.0001).

## ACKNOWLEDGEMENTS

We thank Rachel Hainline for technical assistance with competition assays, Christopher Beierschmitt for technical assistance with worm pathogenicity assays, and Konstanze Schiessl for help with image analysis and feedback on the manuscript.

---

**Figure 2— figure supplement 1.**
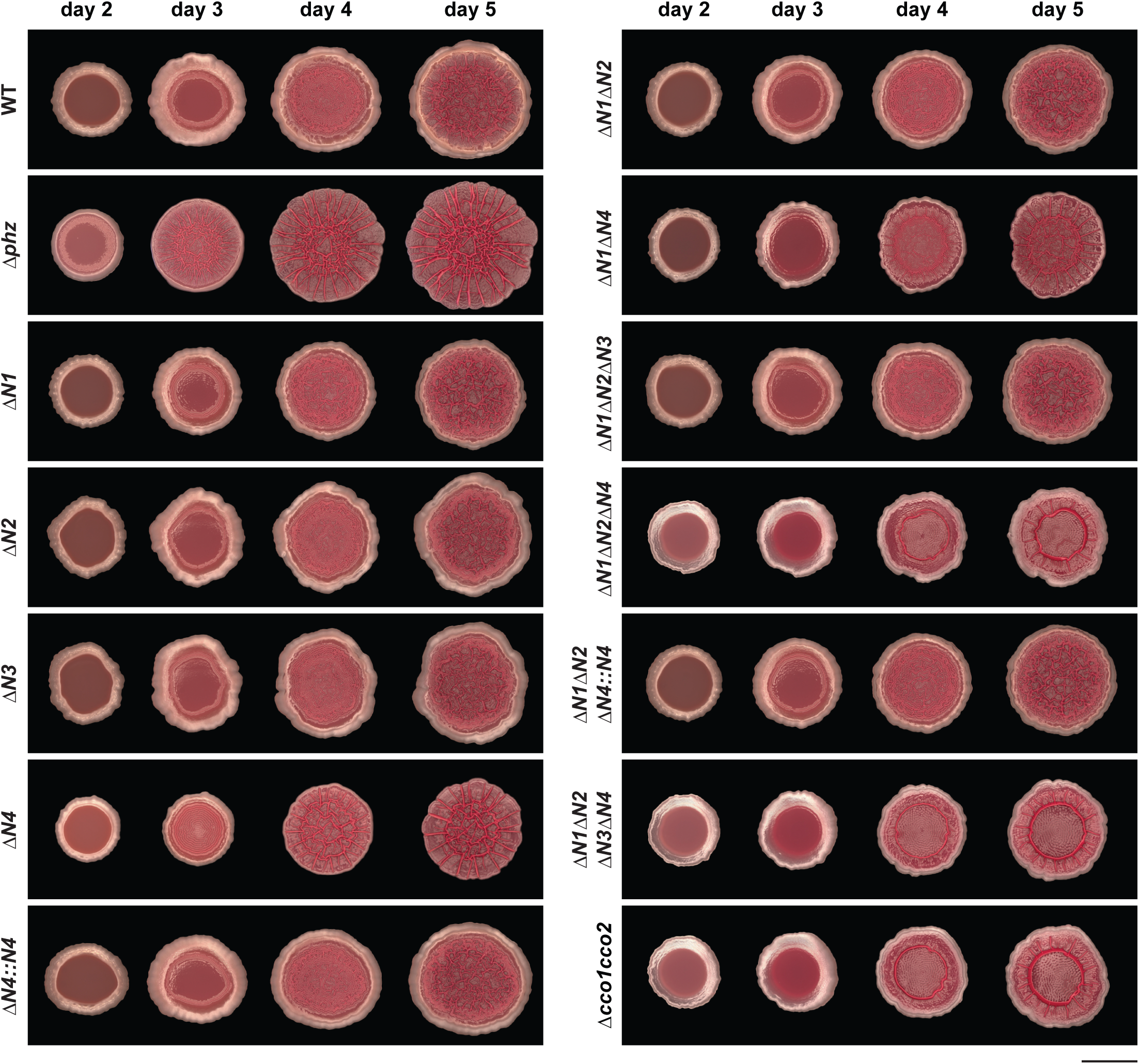
Effects of individual and combined *cco* gene deletions on colony biofilm morphogenesis. Development of WT, Δ*phz*, and *cco* single, combinatorial, and *ccoN4* complementation strains over five days of growth. Images shown are representative of at least five biological replicates and were generated using a Keyence digital microscope. Scale bar is 1 cm.

**Figure 2— figure supplement 2.**
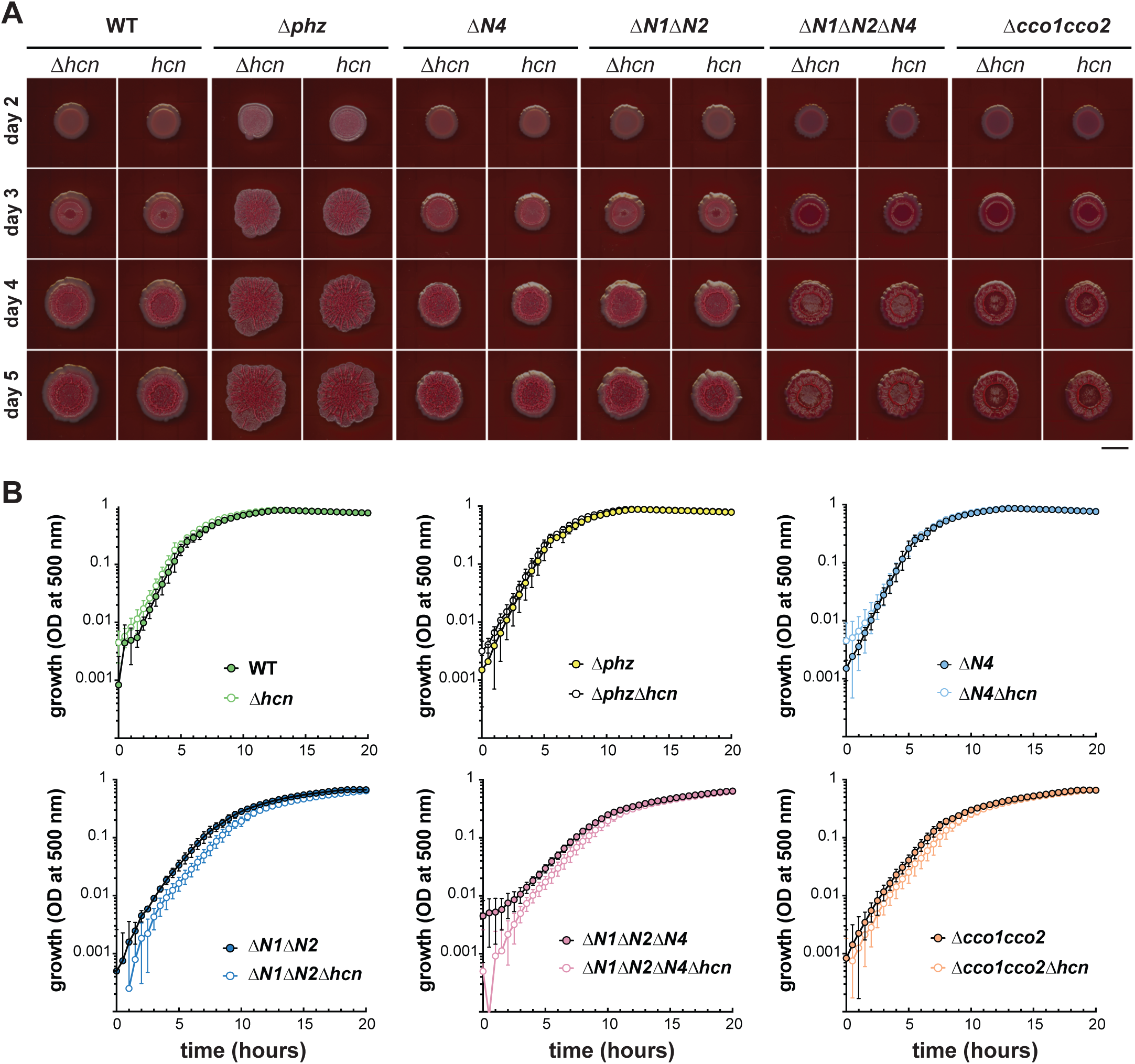
PA14 WT, Δ*phz*, and *cco* mutant colony biofilm phenotypes are unaffected by endogenous cyanide production. (A) Colony development over four days for Δ*phz*, Δ*hcnABC*, and *cco* combinatorial mutants. Images were generated using a flatbed scanner (Epson Expression 11000XL) and are representative of at least three biological replicates. Scale bar is 1 cm. (B) Growth of Δ*phz*, Δ*hcnABC*, and *cco* combinatorial mutants in MOPS defined medium with 20 mM succinate. Error bars represent the standard deviation of biological triplicates and are not shown in cases where they would be obscured by the point marker.

**Figure 2— figure supplement 3.**
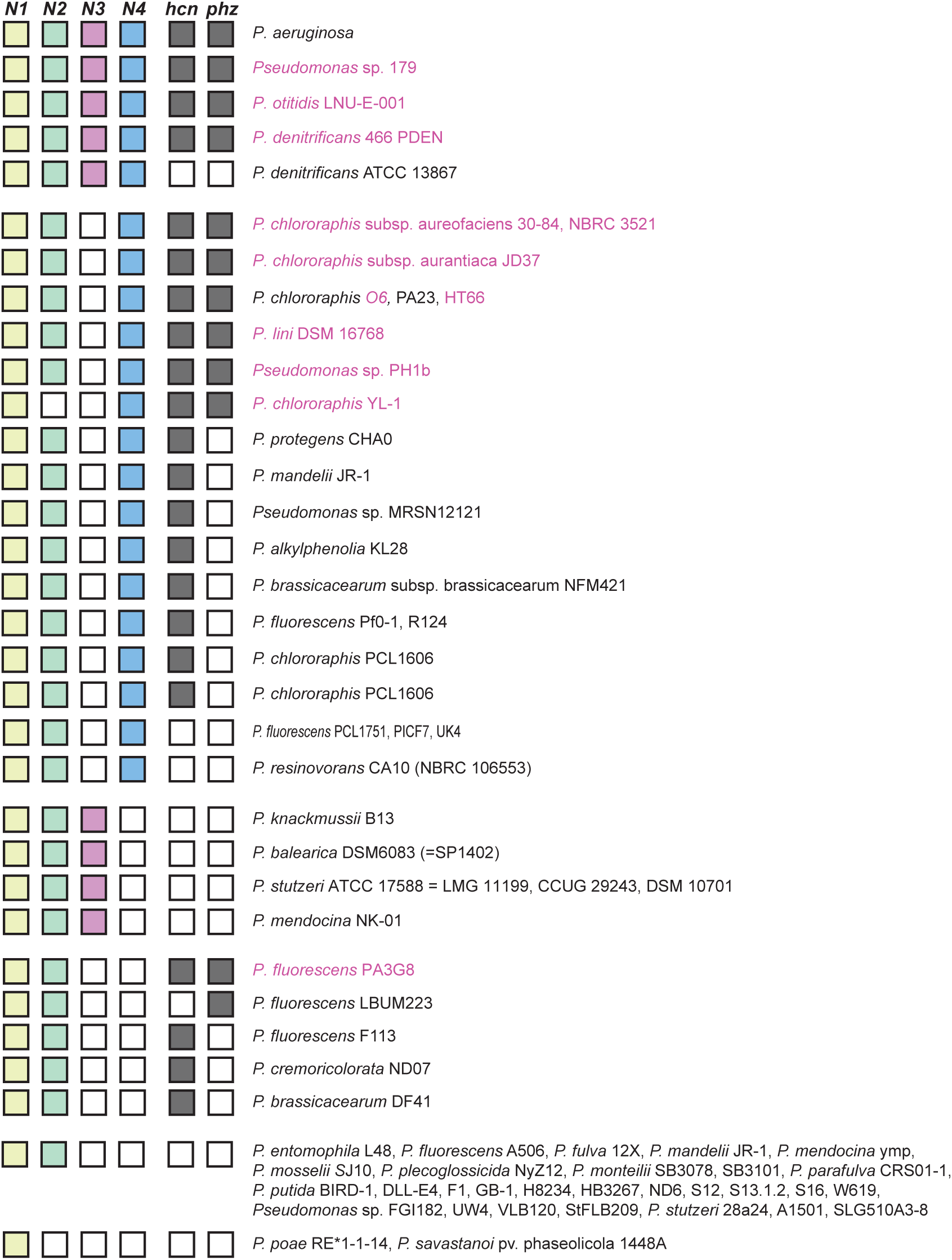
Pseudomonads with CcoN homologs. We examined genomes available in the Pseudomonas Genome Database (Winsor et al. 2016) for CcoN homologs by performing a protein BLAST search on CcoN1 from *P. aeruginosa* PA14. All hits from full genomes, excluding other *P. aeruginosa* strains, were aligned using ClustalW and a tree was built using the geneious tree builder (Geneious 10 (Kearse et al. 2012)). We also included draft genomes (highlighted in purple) that contained genes involved in phenazine biosynthesis (*phzABCDEFG*). The tree revealed four clusters, each being more closely related to one of the four N subunits from PA14, which allowed us to annotate the N subunits accordingly. We next probed all genomes with N subunits for the presence of genes involved in cyanide synthesis (*hcnABC*) and phenazine biosynthesis (*phzABCDEFG*). In contrast to a previous claim, we did not find a clear correlation between the presence of CcoN4 and Hcn proteins (Hirai et al. 2016). We note that with the exception of two *P. fluorescens* strains, those containing *phzABCDEFG* operons also contained *ccoN4.*

**Figure 3— figure supplement 1.**
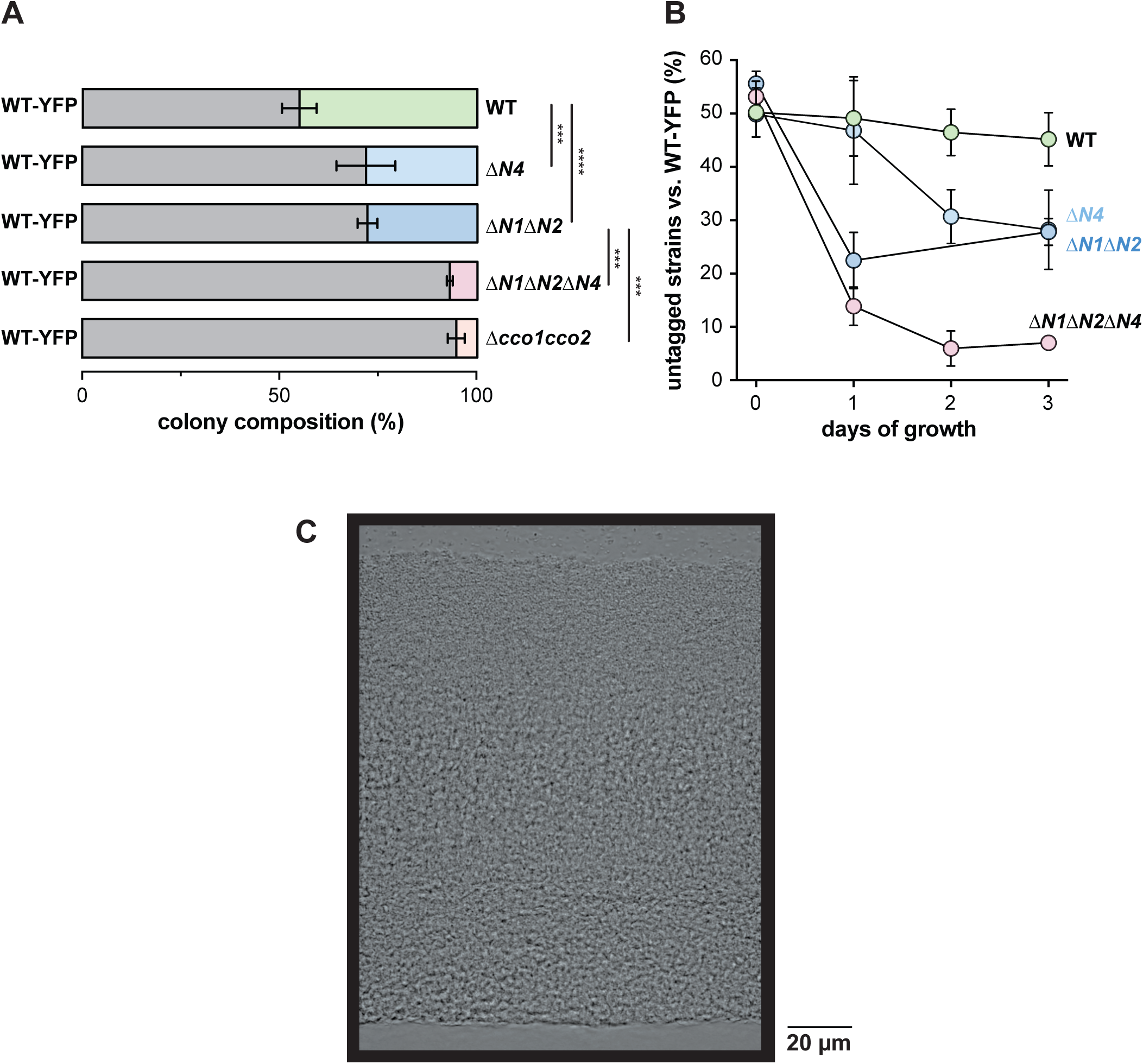
CcoN4 is necessary for optimal fitness in biofilms, particularly when O_2_ becomes limiting. (A) Relative fitness of YFP-labeled WT when co-cultured with various *cco* mutant strains in mixed-strain biofilms for three days. Error bars represent the standard deviation of biological triplicates. P-values were calculated using unpaired, two-tailed t tests (^∗∗∗^, P ≤ 0.001; ^∗∗∗∗^, P ≤ 0.0001). For full statistical reporting, refer to Table 4. (B) Time course showing relative fitness, over a period of three days, of YFP-labeled WT when co-cultured with various *cco* mutant strains in mixed-strain biofilms. Error bars represent the standard deviation of biological triplicates. (C) DIC image of a three-day-old WT biofilm, which is representative of ten biological replicates.

**Figure 4— figure supplement 1.**
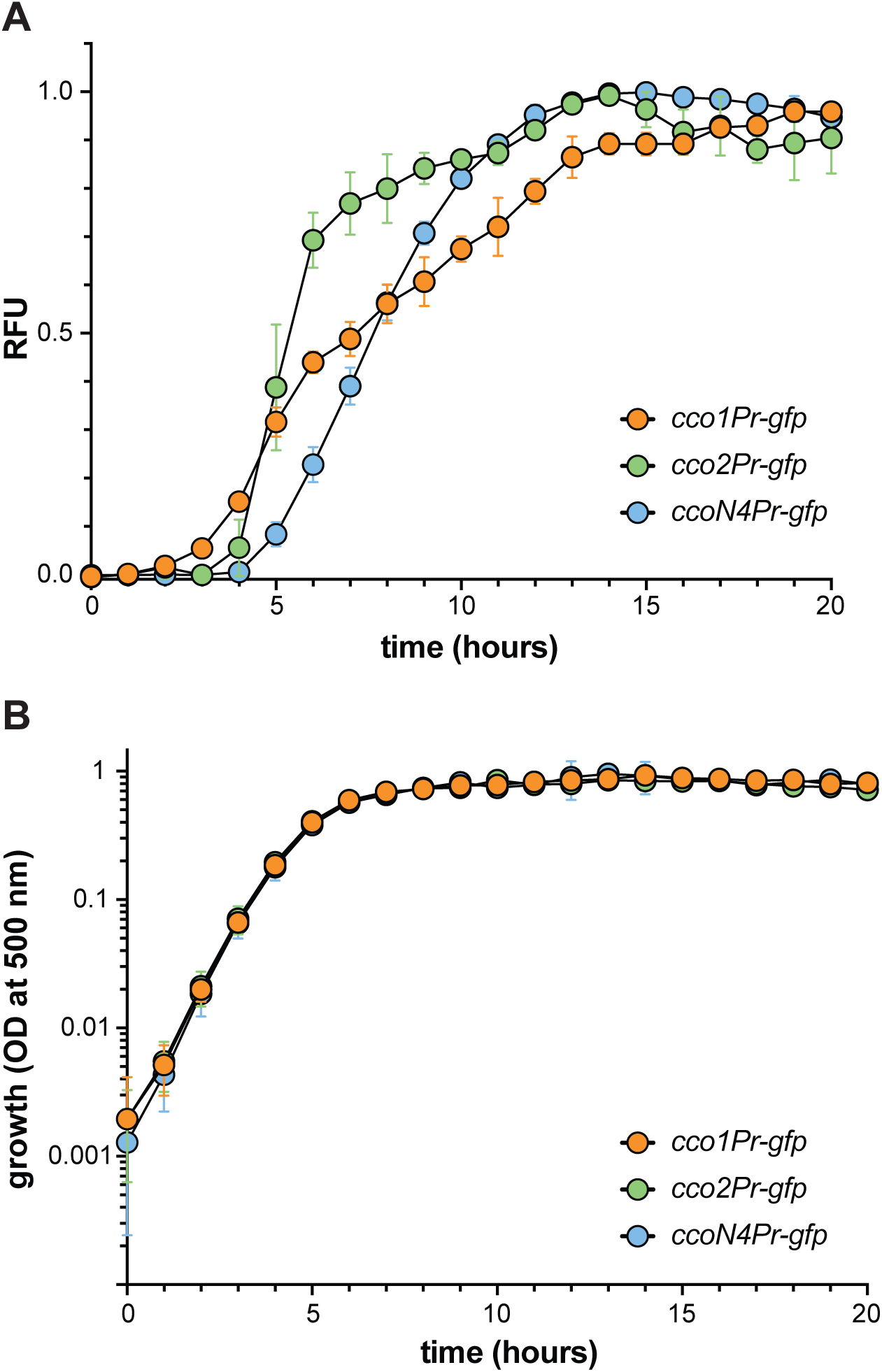
Expression of *cco* reporters in shaken liquid cultures. (A) Fluorescence of translational reporter strains, engineered to express GFP under the control of the *cco1*, *cco2*, or *ccoQ4N4* promoter during growth in 1% tryptone. Fluorescence values for a strain containing the *gfp* gene without a promoter were treated as background and subtracted from each growth curve. (B) Liquid-culture growth of translational reporter strains in 1% tryptone. Error bars in (A) and (B) represent the standard deviation of biological triplicates and are not drawn in cases where they would be obscured by point markers.

**Figure 5— figure supplement 1.**
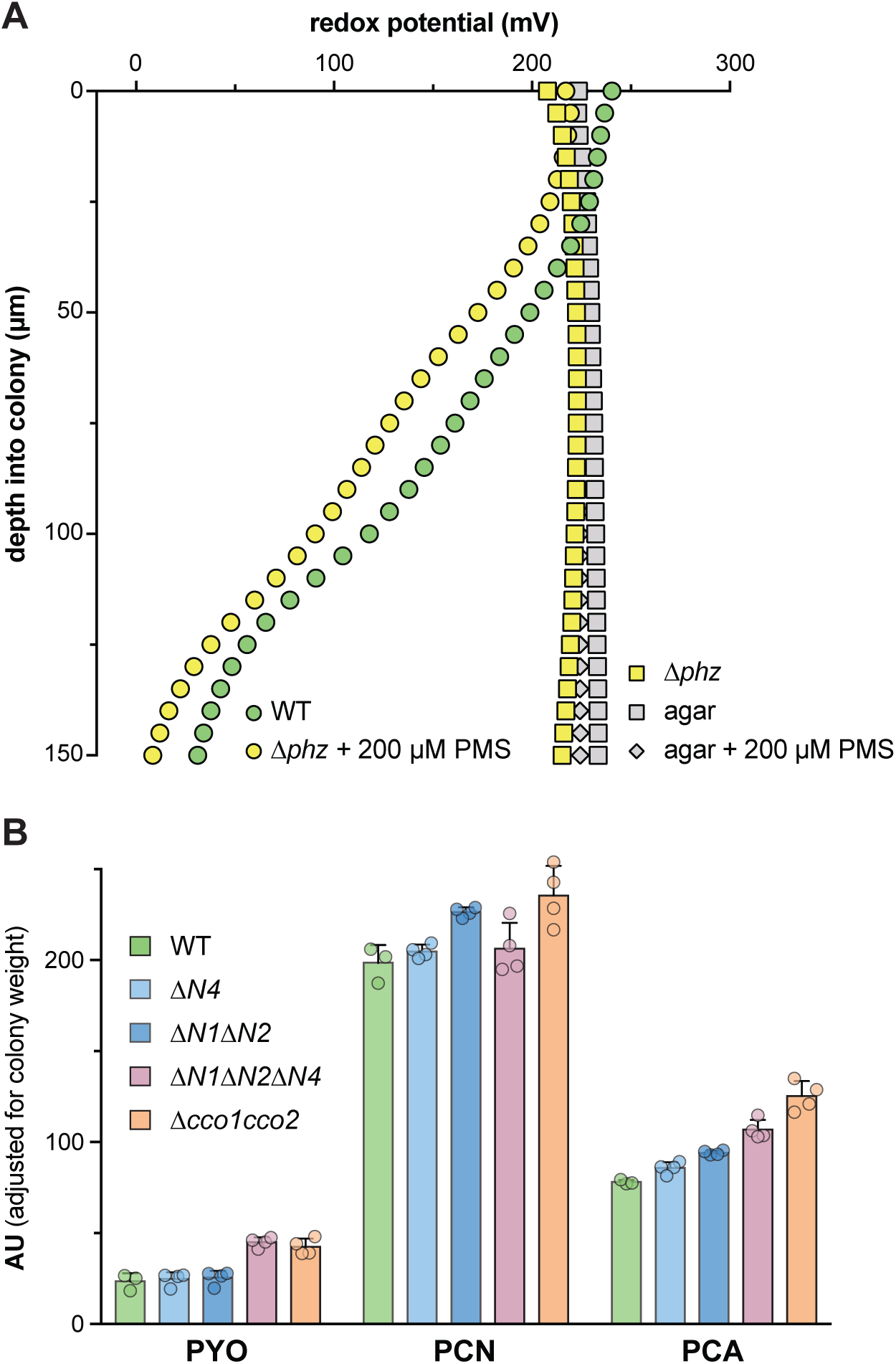
Use of a redox microelectrode to measure phenazine reduction in colony biofilms. (A) Change in redox potential over depth for two-day-old biofilms of PA14 WT, Δ*phz*, and Δ*phz* grown on 200 μM phenazine methosulfate (PMS). Data are representative of at least three biological replicates. To ensure that addition of phenazine methosulfate did not alter the baseline redox potential, a measurement was also taken of agar only. (B) Levels of phenazines extracted from the agar medium underneath the colony and separated by HPLC, adjusted for biomass, for PA14 WT and various *cco* mutant biofilms grown for two days. Data represent the area under each peak in absorbance units for the phenazines indicated, and error bars represent standard deviation of at least three biological replicates.

